# Fluorescence assay for simultaneous quantification of CFTR ion-channel function and plasma membrane proximity

**DOI:** 10.1101/631614

**Authors:** Stella Prins, Emily Langron, Cato Hastings, Emily J. Hill, Andra C. Stefan, Lewis D. Griffin, Paola Vergani

**Author notes:** The first two authors contributed equally to this project.

## Abstract

CFTR, a plasma membrane anion channel, plays a key role in controlling transepithelial fluid movement. Excessive activation results in intestinal fluid loss during secretory diarrhoeas, while *CFTR* mutations underlie cystic fibrosis (CF). Anion permeability depends both on how well CFTR channels work (permeation/gating) and on how many are present at the membrane (reflecting folding, trafficking, metabolic stability). Recently, treatments with two drug classes targeting CFTR – one boosting ion-channel function (potentiators), the other increasing plasma membrane density (correctors) – have provided significant health benefits to CF patients.

Here we present an image-based fluorescence assay that can rapidly and simultaneously estimate both CFTR ion-channel function and the protein’s proximity to the membrane. We monitor F508del-CFTR, the most common CF-causing variant, and confirm rescue by low temperature, CFTR-targeting drugs and second-site revertant mutation R1070W. In addition, we characterize a panel of 62 CF-causing mutations. Our measurements correlate well with published data (electrophysiology and biochemistry), further confirming validity of the assay.

Finally, we profile effects of acute treatment with approved potentiator drug VX-770 on the rare-mutation panel. Mapping the potentiation profile on CFTR structures raises mechanistic hypotheses on drug action, suggesting that VX-770 might allow an open-channel conformation with an alternative arrangement of domain interfaces around site 1.

The assay is a valuable tool for investigation of CFTR molecular mechanisms, allowing accurate inferences on gating/permeation. In addition, by providing a two-dimensional characterization of the CFTR protein, it could better inform development of single-drug and precision therapies addressing the root cause of CF disease.

## Introduction

Anion flow mediated by the cystic fibrosis transmembrane conductance regulator (CFTR), an apical epithelial channel [1], controls volume and composition of the luminal fluid comportment in several organs. CFTR function is thus crucial for physiological processes such as airway mucociliary clearance, secretion of pancreatic juices and maintenance of optimal fluid content in the intestinal lumen [2].

Enterotoxin-induced secretory diarrhoeas are a major global cause of malnutrition, impaired development and death of children [3]. Excessive CFTR-mediated anion conductance (*G*_CFTR_) in the apical membrane of enterocytes causes intestinal loss of large volumes of fluid, leading to dehydration [4]. At the other extreme, cystic fibrosis (CF) a common life-limiting genetic disease [5], is caused by mutations which reduce *G*_CFTR_ throughout the body, severely impacting on life expectation and quality [6,7].

*G*_CFTR_ is the product of 3 factors: the number of channels in the relevant membrane (*N*), channel open probability (*P*_O_), and single-channel conductance (*γ*):

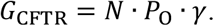

Mutations and bacterial toxins can affect gating and permeation of the mature channel (affecting *P*_O_ and *γ*, respectively). But biogenesis of polytopic CFTR is complex [8,9], and many mutations (and chemical compounds [10]) also impair folding, trafficking and plasma membrane stability, resulting in a smaller number of channels at the membrane (*N*).

Drugs targeting CFTR are emerging: CFTR inhibitors, which could provide emergency treatment for diarrhoeas [11], and CFTR modulators, capable of restoring CFTR activity to defective mutant channels for CF treatment. Modulators belong to two classes: “potentiators” increase *P*_O_, while “correctors” increase plasma membrane density. The potentiator ivacaftor (VX-770, Vertex Pharmaceuticals [12]) dramatically improves lung function of patients carrying G551D [13] or other mutations impairing channel function. Corrector VX-809 [14] is part of new triple combination therapies, combining two different correctors with a potentiator. These have recently brought remarkable clinical benefits to patients carrying at least one copy of the common F508del-CFTR variant, covering ~ 90 % of the CF population [15,16].

Despite these major clinical success stories, little is known on how modulators work. An atomic-resolution structure of a VX-770-bound CFTR [17], reveals the superficial binding of the drug molecule at the interface between transmembrane domain and lipid bilayer. But the binding of the drug is not seen to cause any significant conformational change, (compare VX-770 bound 6O2P [17] vs. 6MSM [18]), and the permeation pathway remains closed [17,18]. How does VX-770 binding increase *P*_O_ of WT-CFTR and many mutant CFTR versions?

To investigate questions such as these and test mechanistic hypotheses, an assay that allows rapid functional screening of changes caused by mutations or compound modification would be useful. But currently available (relatively high throughput) assays report on either CFTR surface expression (e.g. [19,20]) or CFTR-mediated cellular conductance [21]. Apart from low-throughput single-channel patch-clamp recording, assays that measure CFTR function cannot simultaneously measure how many channels are contributing to such function. They cannot discriminate whether a measured conductance arises form a small number of channels with high (*P*_O_ · *γ*) or a larger number of channels with less favourable gating/permeation characteristics.

Here we present a “high-content” assay based on dual-colour live imaging of HEK293 cells, that extracts information on both key characteristics of CFTR: by co-expressing soluble mCherry with the halide sensitive YFP [22] linked to CFTR [23], our new assay gives simultaneous estimates of both CFTR function, and CFTR membrane proximity. Experimental manipulations - incubation at low temperature [24–26], treatment with VX-809 [27,28] with and without VX-770 [29,30], and addition of revertant mutation R1070W [28,31,32] - result in the expected changes in measured F508del-CFTR channel function and membrane proximity. In addition, we present a screening platform suitable for describing the molecular characteristics of 62 missense CFTR variants carried by CF patients, and we profile the effects of VX-770 on this panel. Measurements we obtain correlate well with published datasets, validating our assay as a new tool to investigate questions on CFTR molecular mechanisms and pharmacology.

## Results

### The assay

#### Ion channel function

Expression of a cytosolic halide sensitive YFP with increased affinity for iodide and a low affinity for chloride, YFP(H148Q/I152L) [22,33], allowed the first high throughput CFTR screening projects, assessing CFTR activity by measuring the rate of YFP fluorescence quenching caused by iodide/chloride exchange across the plasma membrane [34–37]. To obtain quantitative information about ion channel function, we fused this YFP to the intracellular N-terminal of CFTR [23,38]. We constructed the pIRES2-mCherry-YFPCFTR plasmid that directs co-expression of YFP(H148Q/I152L)-CFTR (hereafter designated YFP-WT-CFTR or simply WT-CFTR) and a soluble, cytosolic, red fluorescent protein, mCherry [39], with both coding sequences transcribed on a single bicistronic mRNA. HEK293 cells are transiently transfected, and images are automatically acquired (before and after iodide addition) and analysed. The time course of YFP quenching in response to extracellular iodide addition informs on anion conductance. Thanks to the common mRNA, mCherry expression serves as an internal standard for the normalisation of YFP-CFTR expression, reducing variability due to unequal transfection efficiency.

#### Membrane proximity

mCherry expression also allows image segmentation and accurate localization of the cell membrane by marking the border of cells. The “membrane-proximal zone” is defined as comprising a ~1 μm wide band, on the inside of a cell’s boundary (Figure 1A). To obtain a robust relative estimate of the number of channels (*N*) giving rise to the cellular conductance (*G*_CFTR_), we estimate overall “CFTR membrane proximity” in each cell calculating the metric *ρ*. This is obtained by dividing the average YFP-CFTR fluorescence intensity within the membrane-proximal zone (*F*_YFP membrane_), by the average mCherry fluorescence over the entire cell (*F*_mCherry cell_). The *ρ* metric can be thought of as the product of the *F*_YFP membrane_/*F*_YFP cell_ metric, the proportion of YFP-CFTR within the membrane-proximal zone, multiplied by the metabolic stability of YFP-CFTR with respect to mCherry (*F*_YFP cell_/*F*_mCherry cell_). Thus, changes in *ρ* metric will reflect not only changes in efficiency of CFTR maturation and trafficking, but also changes in the overall rates of biosynthesis vs. degradation of the protein.

**Figure 1.**
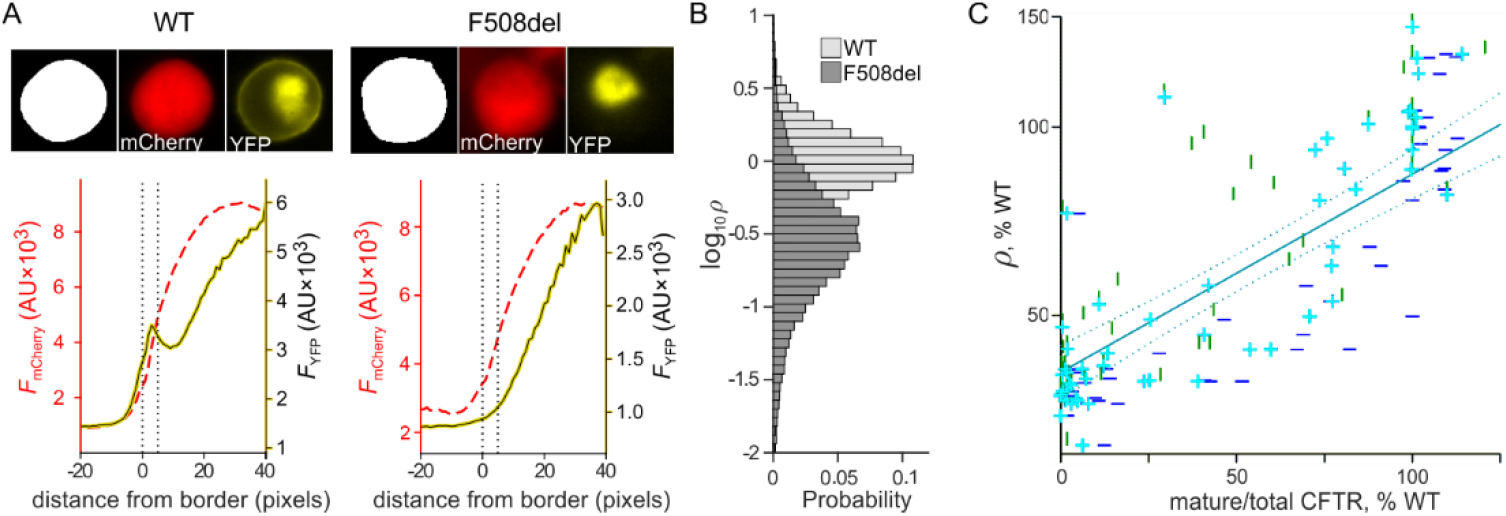
Quantifying CFTR membrane proximity (**A**) Image analysis of individual representative HEK293 cells transfected with pIRES2-mCherry-YFP-WT-CFTR (left), and pIRES2-mCherry-YFP-F508del-CFTR (right). Upper panels: boundary delimiting cell (white) from non-cell (black) is obtained from mCherry image (centre). CFTR cellular localization is obtained from YFP image (right). Lower panels: average mCherry fluorescence intensity (*F*_mCherry_, red dashed line, AU: arbitrary units), and average YFP fluorescence intensity (*F*_YFP_, solid yellow line), as a function of the distance from cell border. Membrane proximity, *ρ*, is defined as

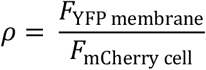

where *F*_YFP membrane_ is the average fluorescence intensity within the ‘membrane proximal zone’, set between 0 and 5 pixels from the cell border (vertical dotted lines). For the representative cells shown WT: *ρ* = 1.60; F508del: *ρ* = 0.25. (**B**) Probability distribution of log_10_*ρ* for cells expressing YFP-WT-CFTR (light grey), and YFP-F508del-CFTR (dark grey), incubated at 37 °C. (**C**) Correlation between the *ρ* metric and published data on complex glycosylation. The latter were obtained from quantifying the ratio (C-band /(C-band + B-band) in Western blots, from FRT cell lines stably expressing missense mutation CFTR variants. Vertical green lines relate our rare-mutations panel with data from [40,42] (*r*^2^=0.53); horizontal blue lines with [41] (*r*^2^=0.74); cyan plus signs with averaged values from the latter two datasets (*r*^2^=0.67). Solid and dotted cyan lines are regression line and 95% confidence intervals, respectively, for the average dataset.

The distribution of *ρ* measurements, easily obtained for hundreds of cells in individual images, is skewed, but approaches a log-normal distribution. Values were log transformed (Figure 1B) before performing statistical analysis.

The *ρ* metric is related to a commonly used measure of CFTR biogenesis, the proportion of protein acquiring complex glycosylation (i.e. that has undergone Golgi processing), estimated using protein blotting. For a set of CF-causing missense mutations (see rare-mutation panel, below), we found a very good correlation (*r*^2^ = 0.67) of our *ρ* measurements with published datasets [40–42] (Figure 1C, see also Supporting Information S8). Note that methodologies and materials used were different: fluorescence measurements in transiently expressing HEK293 cells vs. Western blots from stably expressing Fischer Rat Thyroid, (FRT) cell lines.

For both methodologies, CFTR proteins located in post-Golgi, sub-membrane compartments cannot be discriminated from those at the plasma membrane, directly contributing to *G*_CFTR_. Nevertheless, both measurements, by detecting defects in processing and metabolic stability, provide useful rough estimates of relative plasma membrane numbers.

### Rescue of F508del-CFTR membrane proximity

As a first validation of our assay, we assessed changes in F508del-CFTR membrane proximity by comparing distributions of log_10_*ρ* (logarithmic transformation of the *ρ* metric) following treatments/mutations known to partially rescue the F508del processing defect (Figure 2).

**Figure 2.**
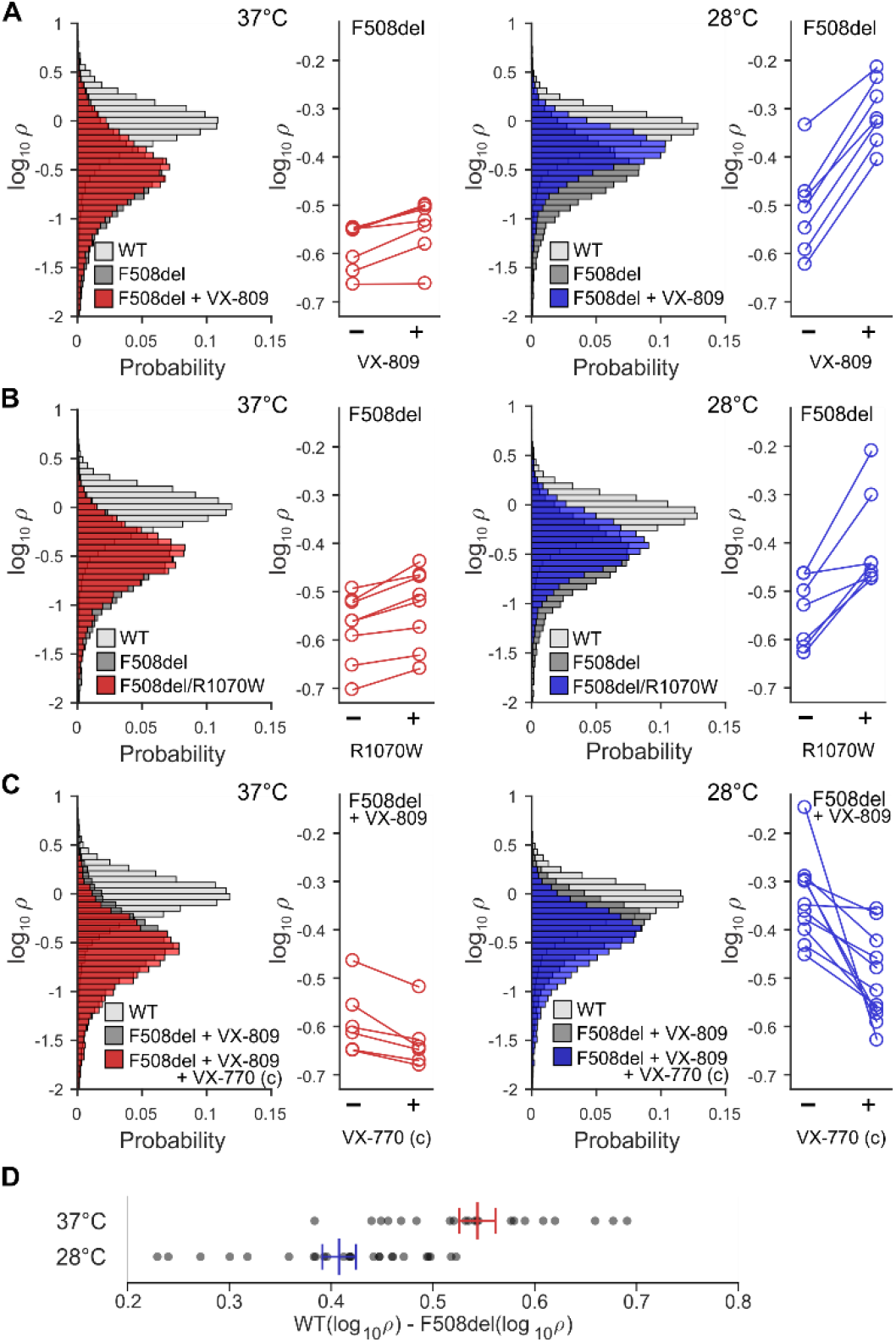
Quantifying rescue of F508del-CFTR membrane proximity. Effects of chronic treatment with 10 μM VX-809 (**A**), R1070W rescue (**B**), and chronic treatment with 10 μM VX-809 ± 10 μM VX-770 (**C**), on log_10_*ρ* at 37 °C (left, red) and 28 °C (right, blue). Conditions of final incubation were maintained during image acquisition. The probability distributions in the panels on the left, contain log_10_*ρ* measurements from thousands of cells, pooled from all experiments. For statistical analysis, mean log_10_*ρ* values determined in independent experiments (individual 96-well plates), and paired per plate, were used (displayed in panels on the right, line connecting measurements from the same plate) (**D**) Before imaging, plates were incubated at 37 °C or 28 °C for 24 hours. For each plate, the difference between mean log_10_*ρ* for WT-CFTR and F508del-CFTR was calculated (WT(log_10_*ρ*) − F508del(log_10_*ρ*), grey dots). Red (37 °C) and blue (28 °C) lines show mean ± SEM, calculated from 21(37 °C) and 25(28 °C) within-plate difference estimates.

#### F508del-CFTR membrane proximity rescue by VX-809 incubation

At 37 °C, incubation with corrector drug VX-809 for 24 hours caused a very small, but significant, increase in log_10_*ρ* of F508del-CFTR, (Figure 2A left, see also Supporting Table S1). At 28 °C, the magnitude of the increase was greater (Figure 2A right).

#### F508del-CFTR membrane proximity rescue by R1070W second-site revertant mutation

Introducing the mutation R1070W, known to partially revert the F508del-CFTR defective phenotype [32], significantly increased F508del-CFTR membrane proximity at 37 °C (Figure 2B left, Supporting Table S1), as well as at 28 °C (Figure 2B right, Supporting Table S1). Again, the magnitude of the effect was larger at 28 °C.

#### F508del-CFTR membrane proximity decrease due to chronic VX-770 incubation

When comparing cells expressing F508del-CFTR incubated for 24 hours with VX-809 alone, with those incubated with both corrector VX-809 and potentiator VX-770, at 37 °C, there was a small but significant decrease in log_10_*ρ* (Figure 2C left, Supporting Table S1). At 28 °C the decrease was again more pronounced than at 37 °C (Figure 2C right).

#### F508del-CFTR membrane proximity rescue by temperature correction

Temperature could only be varied between plates, preventing the use of within-plate differences in log_10_*ρ* to directly compare membrane proximity of F508del-CFTR incubated at different temperatures. We therefore compared the magnitude of the within-plate difference between F508del-CFTR and WT-CFTR for plates incubated at 28 °C and at 37 °C. The log_10_*ρ* values of F508del-CFTR were significantly closer to those of WT-CFTR at 28 °C than at 37 °C, (Figure 2D, Supporting Table S1).

### Rescue of F508del-CFTR ion channel function

Functional rescue of F508del-CFTR was also measured. In these experiments, CFTR was activated following addition of extracellular I^−^ (I^−^ first Protocol, see Experimental Procedures). Activation occurred either by addition of only 10 μM forskolin, increasing intracellular cAMP, and thus CFTR phosphorylation, or by addition of a combination of 10 μM forskolin and 10 μM VX-770 (the latter defined as an acute (a) treatment, as opposed to the 24-hour chronic (c) incubation with VX-770 described above). Normalized YFP fluorescence was followed over time (Figure 3). The maximal rate of I^−^ entry (*Δ*[*I*^−^]_in_/*Δt*) was used to summarize CFTR channel function for the different CFTR genotypes, incubation and activation conditions tested (Figure 3E, Supporting Tables S2 and S3). No significant difference in this metric was detected among the different genotypes/conditions when DMSO (vehicle) was added instead of activators.

**Figure 3.**
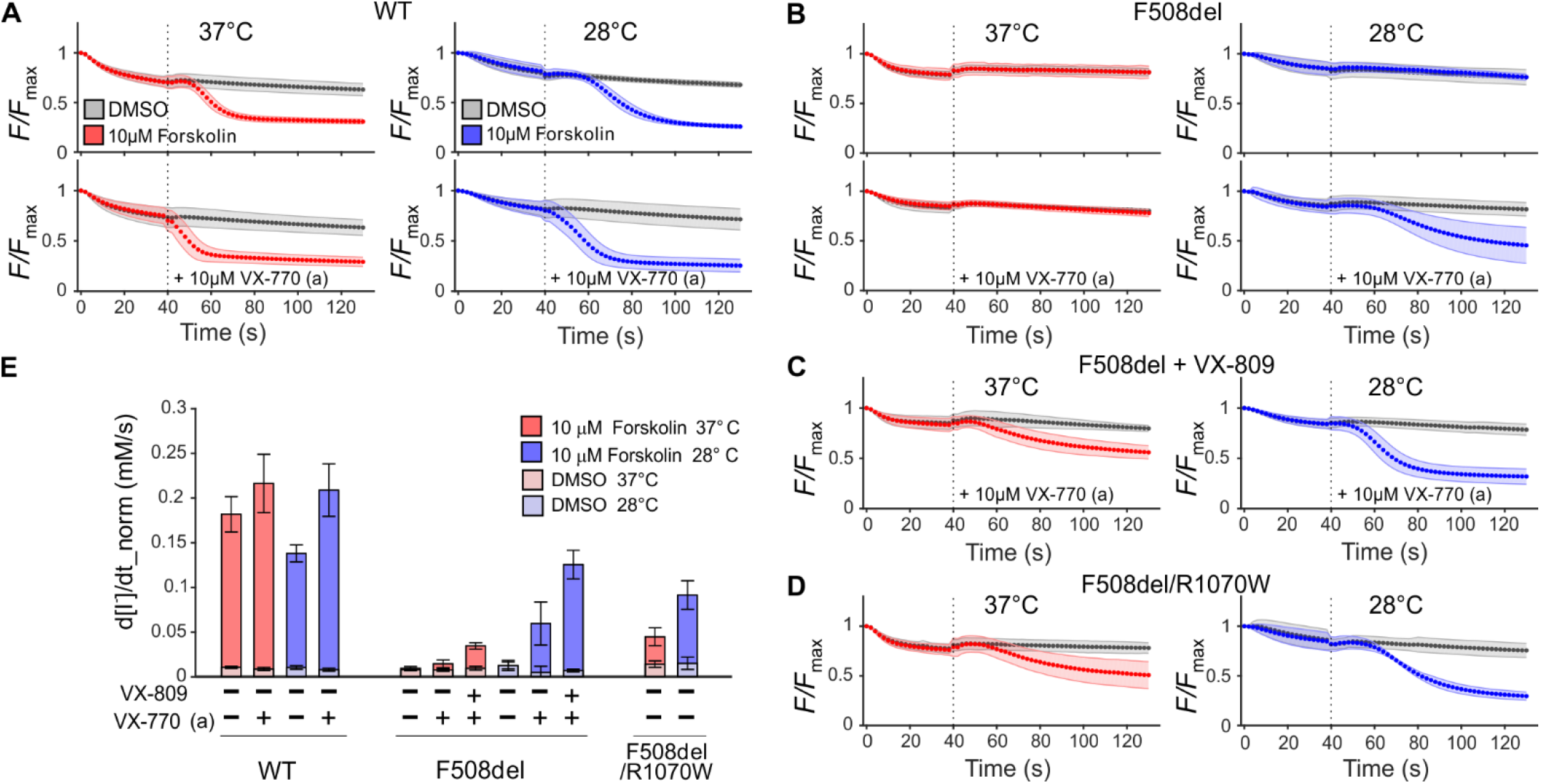
Rescue of F508del-CFTR ion channel function. (**A-D**) Quenching of YFP fluorescence in HEK293 cells expressing WT-CFTR (**A**), expressing F508del-CFTR chronically (24 h) treated with vehicle only, DMSO (**B**), or with VX-809 (**C**), or expressing R1070W/F508del-CFTR (DMSO only chronic treatment) (**D**). *F*/*F*_max_: observed YFP fluorescence, normalized using fluorescence at the time point before I^−^ addition. For more information on statistical analysis see Supporting Tables S2 and S3. Prior to imaging plates were incubated for 24 hours, at 37 °C (red) or 28 °C (blue). This final incubation temperature was maintained throughout image acquisition. At time point 0 s I^−^ was added to the extracellular medium. At 40 s (dotted line) forskolin and, where indicated, VX-770 (acute, a) was added, both to a final concentration of 10 μM. (**E**) The maximal rate of I^−^ entry (d[I^−^]/dt_norm) is used to summarize CFTR function for genotypes and conditions shown in (**A**-**D**).

#### WT-CFTR

Measurements from HEK293 cells expressing WT-CFTR were taken for comparison purposes. As expected, the maximal rate of I^−^ entry was significantly higher after activation with forskolin, compared to control (DMSO), at both 37 °C and 28 °C (Figure 3A; Figure 3E WT). However, conditions were optimised for measuring low CFTR activity, and neither the presence of 10 μM VX-770 in addition to forskolin during activation, nor incubation at 37 °C vs. 28 °C increased quenching rate sufficiently to achieve statistical significance after multiple comparison correction (Figure 3A; Figure 3E, WT, Supporting Table S3).

#### F508del-CFTR functional rescue following temperature correction

Activation with forskolin alone failed to increase the maximal rate of I^−^ entry in untreated cells expressing F508del-CFTR (Figure 3B top; Figure 3E F508del bars 1 and 4, Supporting Table S2), reflecting the severe gating defect, which persists even after temperature correction. Acute potentiation by VX-770 was required to detect function of the channels reaching the cell surface thanks to temperature correction (Figure 3B, bottom; Figure 3E F508del bars 5 vs. 2, Supporting Table S2).

#### F508del-CFTR functional rescue following VX-809 correction

At both temperatures, acute potentiation revealed the activity of F508del-CFTR channels that had reached the cell surface thanks to 24-hour incubation with VX-809. At 28 °C the maximal rate of I^−^ entry was significantly greater than at 37 °C (Figure 3C; Figure 3E, F508del bar 6 vs. 3, Supporting Table S3).

#### F508del-CFTR functional rescue by the R1070W mutation

Forskolin activation alone was enough to reveal F508del/R1070W-CFTR channel activity (Figure 3D, Supporting Table S2). The maximal rate of I^−^ entry was significantly higher at 28 °C than at 37°C (Figure 3D; Figure 3E F508del/R1070W, Supporting Table S3).

### The rare-mutation panel

More than 300 CF-causing mutations have been characterized (The Clinical and Functional TRanslation of CFTR (CFTR2); available at http://cftr2.org). CF-causing missense CFTR mutations [40–42] were individually introduced in the pIRES2-mCherry-YFPCFTR plasmid, creating a panel of 62 plasmids (including WT-CFTR as reference).

Following expression of the panel in HEK293 cells, and incubation with no pharmacological correction, distributions for the *ρ* metric, and plate log_10_*ρ* means were obtained (Supporting Table S4, Supporting Figure S5). The data is summarized in Figure 4A, which profiles membrane proximity for each CFTR mutant variant in the panel.

**Figure 4.**
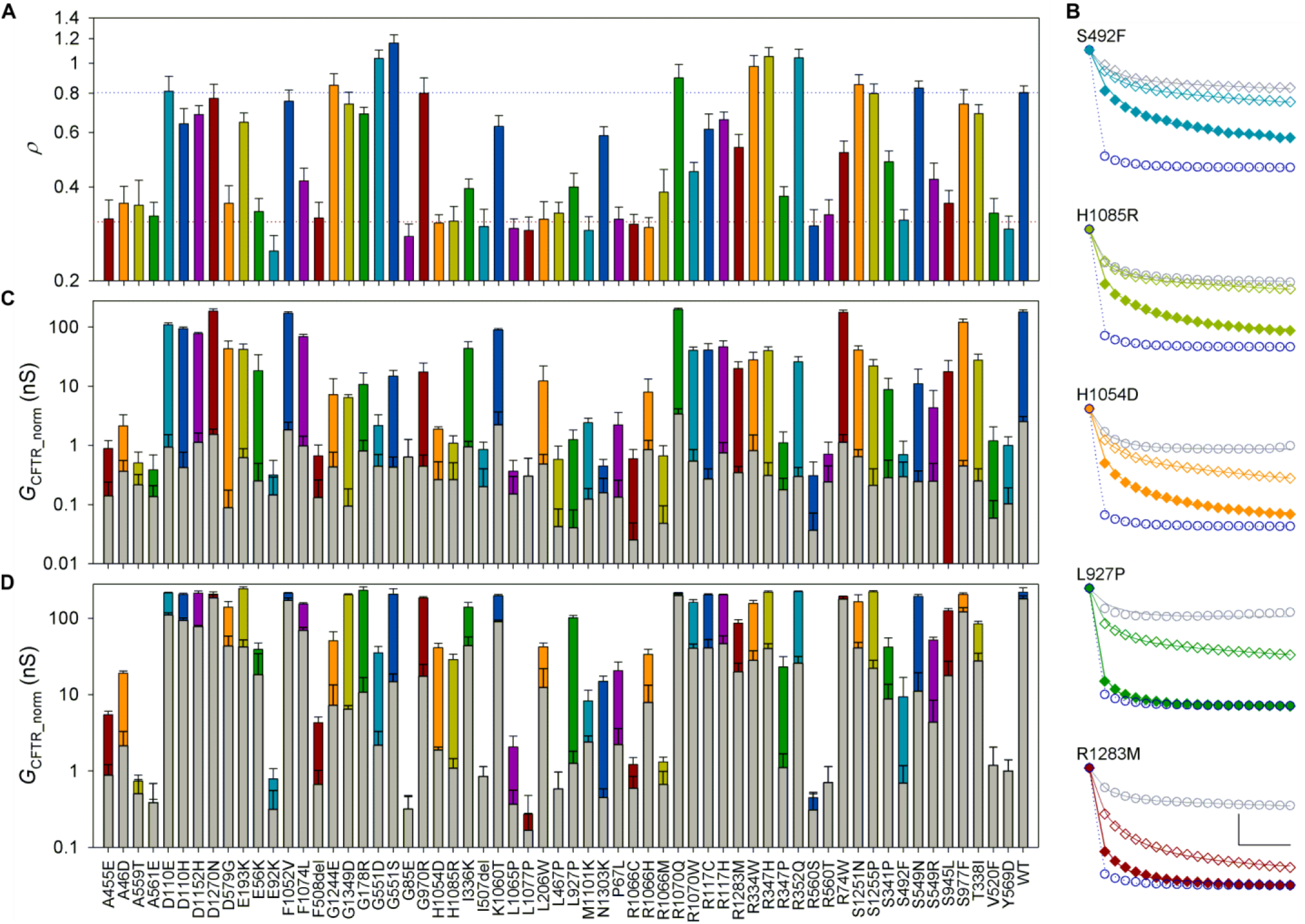
Rare CF-mutation profiling. (**A**) Mean *ρ* (n ≥ 9) of all mutations in the panel. Blue and red dotted lines indicate mean *ρ* for WT- and F508del-CFTR, respectively. For *ρ* distributions, mean *ρ* and n values for each mutant see Supporting Figure S5 and Supporting Table S4. (**B**) Observed YFP fluorescence quenching time course after activation with DMSO (grey circles), or 10 μM forskolin (empty coloured diamonds), or 10 μM forskolin + 10 μM VX-770 (a) (filled coloured diamonds) for selected mutations. Solid lines show predicted change in proportion of anion-free YFP. For estimated parameters *G*_CFTR_, *V*_M_, *G*_trans_ and τ_trans_ see Supporting Table S9. WT-CFTR quenching in 10 μM forskolin (dark blue empty circles, observed, and dotted line, fit) shown for comparison. (**C**) CFTR conductance of rare-mutation panel after activation with 10 μM forskolin (coloured bars) or vehicle control (DMSO, grey bars). n ≥ 3. *G*_CFTR_ obtained from fitting of quenching time-course for each mutant was normalized using the mean within cell mCherry fluorescence for that mutant, measured with respect to the corresponding metric obtained for WT-CFTR on the same plate. For statistical analysis see Supporting Table S6. (**D**) Potentiation of rare-mutation panel by VX-770. Grey bars show values following activation with 10 μM forskolin alone, coloured bars with further addition of 10 μM VX-770 (a). For statistical analysis see Supporting Table S7.

As mentioned above, correlation between our measured *ρ* and the proportion of CFTR protein acquiring complex glycosylation in FRT cells is very good (*r*^2^ = 0.74 [41], *r*^2^ = 0.53 [40,42], and *r*^2^ = 0.67 using average values for mutants measured by both groups [40–42], Figure 1C and Supporting Figure S8).

Time course of YFP fluorescence quenching was also acquired and analysed (I^−^ last Protocol, see Experimental Procedures). In these experiments, steady-state CFTR conductance (*G*_CFTR_) was estimated, with no activation (DMSO) or following baseline pre-activation with 10 μM forskolin (Figure 4B-C; Supporting Table S6). Again, results correlate well with published data (*r*^2^ = 0.68 [41], *r*^2^ = 0.61 [40,42], *r*^2^ = 0.60 [40–42], Supporting Figure S8). Conductance was also measured following activation with 10 μM forskolin + 10 μM VX-770 (a) (Figure 4B, D; Supporting Table S7). In these conditions, genotypes with high conductance (including WT-CFTR) have faster YFP quenching than can be reliably measured in our system. However, the assay can accurately monitor VX-770 potentiation when CFTR activity is low, as is the case for most mutants.

### Relationship between CFTR ion channel function and membrane proximity

By considering changes in ion channel function in the context of any change measured in *ρ*, our assay allows accurate inferences on the gating and permeation properties of the CFTR channel molecules present at the cell surface.

Even when virtually no channels are present in the plasma membrane (as happens, for instance, for cells expressing F508del-CFTR grown at 37° C) the value of *ρ* does not fall to zero. This is likely due to some inaccuracy in automated cell boundary delineation and to the widefield microscope optics, resulting in stray light from out-of-focus planes reaching the photomultiplier. To empirically investigate the relationship between *G*_CFTR_ and *ρ*, cells expressing F508del-CFTR were treated with increasing concentrations of corrector VX-809, progressively improving both biogenesis/membrane stability and conductance (Figure 5A-B). Measured *G*_CFTR_ values as a function of *ρ* values show a roughly linear relationship (Figure 5B, dotted green line). The line can be extended to cross the *ρ* axis, extrapolating to an intercept at *ρ* = 0.23. In addition, in as much as *ρ* values are proportional to the number of channels at the membrane (*N*), the steepness of this line is an estimate of the product (*P*_O_·*γ*). An extension of the line towards higher membrane proximity values shows the *G*_CFTR_ values expected with a higher number of channels reaching the membrane, but retaining gating/permeation characteristics of F508del-CFTR, acutely potentiated by VX-770. It can be seen that, in these conditions, F508del-CFTR is characterised by *P*_O_ levels similar to those of WT-CFTR (the latter without potentiation, Figure 5B, large dark blue empty circle, not far above dotted green line), consistent with patch-clamp measurements (note that *γ* is unaffected by the F508del mutation) [43,44].

**Figure 5.**
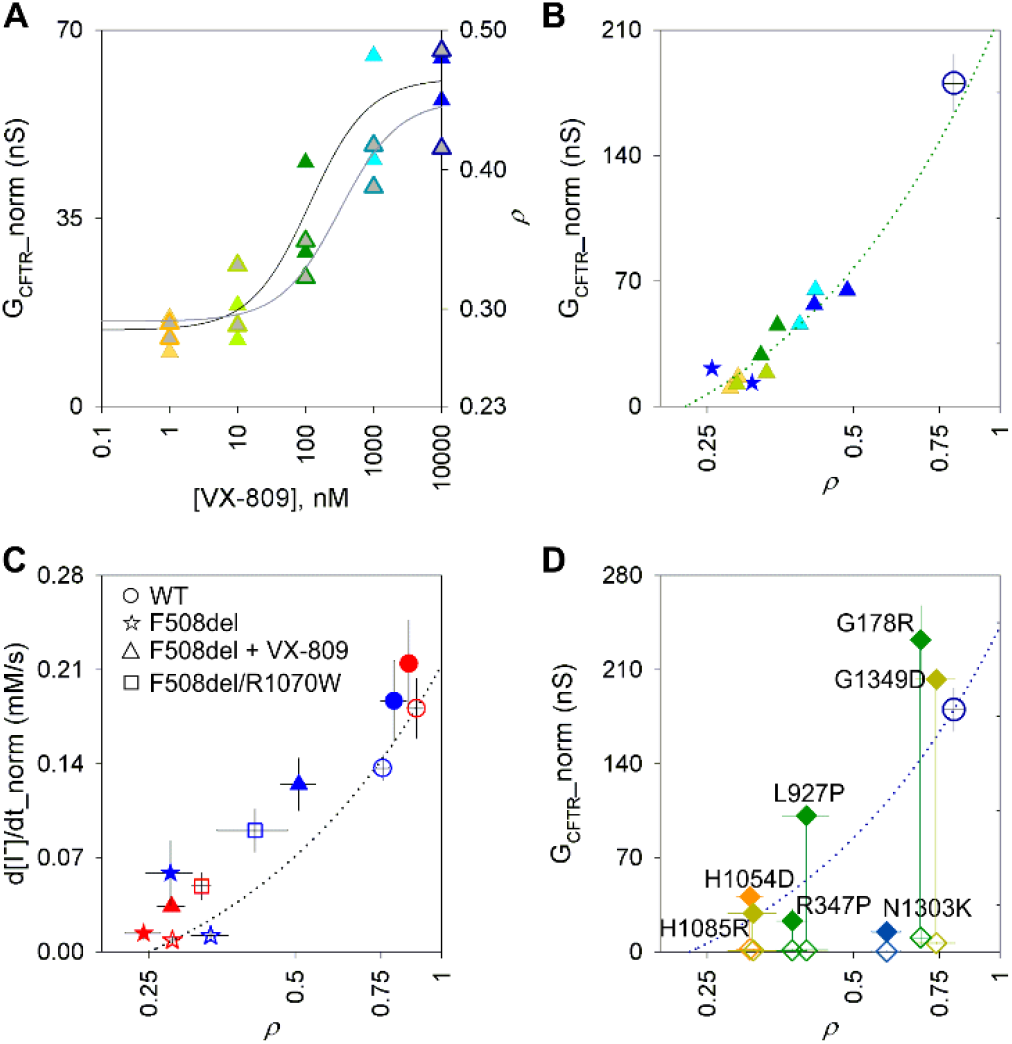
Inferring permeation/gating characteristics. (**A**) Dose-response plot of increase in conductance (left axis, coloured symbols, black fit line) and membrane proximity (right axis, grey-filled symbols, grey fit line) following incubation of F508del-CFTR with increasing concentrations of VX-809. Lines represent fits to the Hill equation (4 parameters, n_H_ constrained to 1, see [23]). Only two measurements were taken at each concentration, but the EC_50_ values we obtain (114 nM ± 66 nM and 316 nM ± 238 nM, for G_CFTR_ and *ρ*, respectively) are not dissimilar from published values [14,20]. (**B**) Relationship between normalized CFTR conductance and membrane proximity in cells expressing F508del-CFTR with no correction (blue stars) or incubated with increasing concentrations of VX-809 (1 nM to 10 μM, colour-coded as in panel A), all after activation with 10 μM forskolin and 10 μM VX-770 (a). F508del-CFTR incubation and measurements were at 28 °C. Green dotted line shows linear regression using only F508del-CFTR data points on graph (slope = 281.7, constant = −63.7, resulting in an x-axis intercept at *ρ* = 0.23). Mean value for WT-CFTR activated with 10 μM forskolin alone, large dark blue empty circle, is shown for reference (from (**D**), see also Figure 6). (**C**) Relationship between maximal rate of I^−^ influx and *ρ* in HEK293 cells expressing WT-CFTR, F508del-CFTR, and F508del/R1070W-CFTR, at 37 °C (red symbols) and 28 °C (blue symbols). Empty symbols indicate CFTR activation with 10 μM forskolin alone; solid symbols indicate further acute potentiation with 10 μM VX-770. Dotted line: linear interpolation between data obtained at 37 °C for uncorrected F508del-CFTR (used as an empirical measure of minimal membrane proximity) and WT-CFTR, both without acute VX-770 potentiation; slope = 0.284, constant = −0.071, resulting in an x-axis intercept at *ρ* = 0.25. (**D**) Mutants with largest fold potentiation by VX-770 (ratio between conductance obtained in 10 μM forskolin + 10 μM VX-770 (a) over that in 10 μM forskolin alone > 20). Empty diamonds indicate baseline activation with 10 μM forskolin alone, solid diamonds indicate activation following acute potentiation with 10 μM forskolin + 10 μM VX-770 (a).

Data on maximum rate of I^−^ entry can also be plotted against the corresponding *ρ* values, measured for the different F508del-CFTR rescue strategies (Figure 5C). A linear interpolation between data points for uncorrected F508del-CFTR at 37° C (representing cells with virtually no CFTR molecules at the membrane) and WT-CFTR activated by 10 μM forskolin at 37°C describes the ion channel function we would expect from cells with increasing CFTR membrane proximity, assuming gating and permeation characteristics of baseline-activated WT-CFTR (Figure 5C, blue dotted line). This allows us to infer how the rescued F508del-CFTR channels reaching the membrane compare to control channels in terms of permeation/gating.

Introducing the R1070W revertant mutation in the F508del-CFTR background is shown to be particularly effective in improving gating (note that permeation and single-channel conductance, are unaffected by both F508del and R1070W mutations [32,45]). R1070W revertant rescue and temperature correction similarly increase membrane proximity. However, temperature-corrected F508del-CFTR channels at the membrane have very low ion channel function (unless acutely potentiated with VX-770). In contrast, F508del/R1070W channels at the membrane have gating and permeation properties equal – or even superior – to WT-CFTR (Figure 5C, cf. uncorrected F508del-CFTR blue symbol vs. F508del/R1070W-CFTR red symbol both compared to blue dotted line). Both results are consistent with patch-clamp records indicating a F508del/R1070W-CFTR *P*_O_ comparable to that of WT-CFTR [46], but a much lower *P*_O_ for temperature-corrected F508del-CFTR [43,44,46].

Figure 6 plots *G*_CFTR_ as a function of *ρ* for the rare-mutation panel, giving an immediate representation of how severe a defect each mutation causes in biogenesis (distance from WT-CFTR on the x-axis) and/or in gating and permeation properties (vertical displacement from blue dotted line, which assumes ion-channel properties of baseline-activated WT-CFTR). For instance, D579G-CFTR (orange open diamond at coordinates (0.35,41.5)) falls close to the WT-CFTR line, suggesting that the product *P*_O_·*γ* is not greatly affected by this mutation, and that the low short-circuit currents measured in FRT cells [40,41] are largely caused by the reduced membrane density. For G1244E (orange (0.75,7.2)) and S549N (blue (0.83,11)), likely altering the structure of CFTR’s canonical ATP binding site 2 (in P-loop and signature sequence loop, respectively), measured ion channel function is much lower than would be expected given the high observed membrane proximity. Here low short-circuit currents [41] are likely due to gating defects. Most mutations give reduced membrane proximity and a conductance that falls below the WT interpolation line, suggesting processing defects as well as some degree of impairment in gating/permeation for the CFTR molecules that do reach the membrane. We further illustrate the effect of acute treatment with VX-770 for mutations resulting in the strongest potentiation (fold-potentiation >20, Figure 5D). For many of these, data points for potentiated conductance fall above the interpolation line, suggesting that the product (*P*_O_·*γ*) is higher than measured for WT-CFTR in baseline-activated conditions.

**Figure 6.**
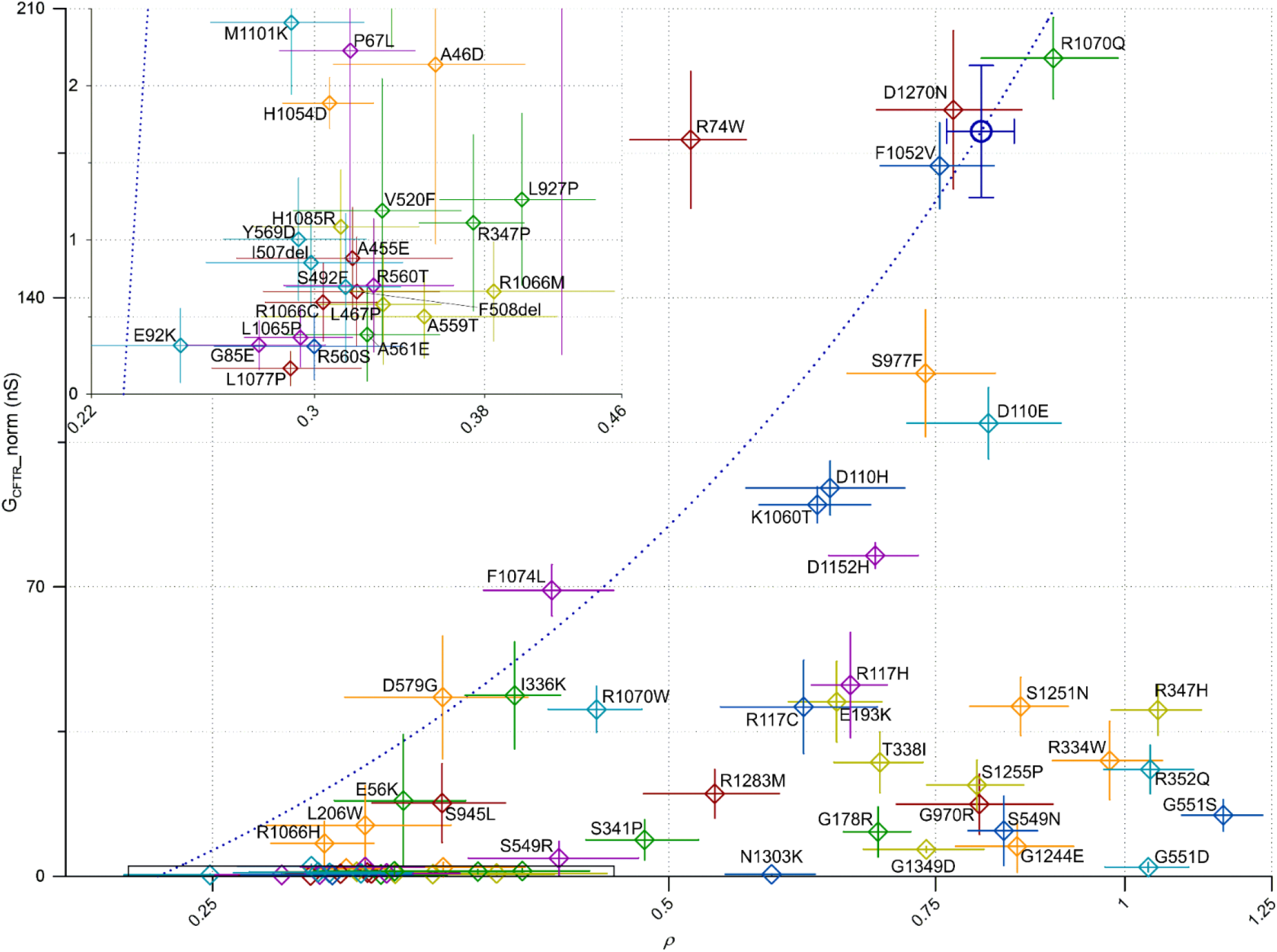
Relationship between baseline *G*_CFTR__norm (10 μM forskolin) and *ρ* for rare-mutation panel. Colours as in Figure 4. WT-CFTR is highlighted as a large, dark blue, empty circle. The dark blue dotted line (slope = 314.1, constant= −72.3) shows linear interpolation between WT data point and x-axis intercept set at *ρ* =0.23, as obtained in Figure 5B. Inset shows expanded axes view of area indicated by black rectangular outline (0 < G_CFTR_norm_ < 2.5 nS; 0.22 < *ρ* < 0.46).

## Discussion

### Validation of the assay

#### Validation of membrane proximity measurements

Although heterogeneity among *ρ* values for individual cells is large, resulting in broad distributions (Figure 2), much of the variability is related to between-plate variation, such that paired comparisons between measurements obtained in the same plate (right panels in Figure 2) can pick up small changes in membrane proximity, increasing assay sensitivity.

For instance, we measure small changes in F508del-CFTR membrane proximity due to incubation with corrector VX-809 at 37 °C. While one published paper reports large effects of this corrector, resulting in rescue of up to 15% of WT-CFTR function [14], much more limited effects are measured by other groups (a 3-4 fold increase in plasma membrane density or function, starting from a value of approximately 1% of WT [28,47]). Our assay detects a change in membrane proximity of a similar magnitude to the latter reports (cf. [28,47] vs. Figure 2A left). These limited *in vitro* effects are more in agreement with the inability of VX-809 monotherapy to improve lung function for F508del homozygous patients [48].

The effect we measure for the R1070W mutation at 37 °C is similarly small, but also significant (Figure 2B left). Again, our result confirms observations published by others: the rescue of membrane-exposed F508del-CFTR due to the R1070W mutation is limited (from 2% to 7% of WT-CFTR), becoming more obvious only when combined with other rescue manoeuvres such as additional revertant mutations or correctors [28].

We could also confirm previous reports demonstrating increased membrane proximity of F508del-CFTR due to low temperature incubation [24–26] (Figure 2D) and enhanced effects of VX-809 treatment when combined with incubation at low temperature [27] (Figure 2A right). We further demonstrate that low temperature incubation also enhances R1070W rescue. The synergy between effects of low-temperature and the R1070W mutation, and of low temperature and VX-809 incubation, suggests that, while VX-809 and the R1070W mutation are acting via a common mechanism stabilizing the NBD1/TMD interface (between nucleotide binding domain 1 and transmembrane domain) [28], a different pathway, possibly involving proteostasis components [26], likely underlies rescue by low-temperature incubation.

In agreement with other studies [29,30,49], we observed a small but significant shift in log_10_*ρ* following chronic incubation with VX-770, consistent with the potentiator destabilizing F508del-CFTR at the membrane (Figure 2C left). Furthermore, we find that the negative effect of VX-770 on biogenesis appears more pronounced when cells are incubated at 28°C (Figure 2C). It is possible that binding of VX-770 prevents interaction with chaperone(s) which help F508del-CFTR fold and exit the ER in cells grown at low temperature [26]. However, the concentration of VX-770 we used (10 μM) is relatively high [50]. Despite the fact that in our incubation medium, as in plasma, a large proportion of the drug will be bound to proteins present in the added serum [51], VX-770 will likely accumulate in the hydrophobic membranes [50,51]. Hence it is also possible that some of the F508del-CFTR destabilization we observe might be linked to formation of precipitates within cellular membranes [50], which would be more pronounced at the lower temperature.

#### The HEK293 expression system

We implemented our assay in the HEK293 heterologous expression system, characterized by robustness, ease of culture and of genetic manipulation. While HEK293 cells do not form monolayers suitable for functional measurements of transepithelial currents, they are widely used in the study of both CFTR function and biogenesis [52–57]. Our measurements of temperature-, VX-809-, and R1070W-dependent recue of F508del-CFTR membrane proximity (Figure 2), confirm results obtained using other systems including human bronchial epithelia [28,47]. In addition, our membrane proximity measurements for the rare-mutation panel (Figure 4A) correlate well (Figure 1C, Supporting Figure S8) with immunoblot measurements obtained with FRT cell lines stably expressing CFTR variants [40,41], a system known to have *in vivo* predictive value for CF [41,47]. Our study thus validates the use of HEK293 cells as a tool for the molecular characterization of the CFTR protein, including its biogenesis.

However, while acute potentiator action is largely independent of the cell system used for testing (e.g. VX-770 is effective in a range of expression systems, from *X.laevis* oocytes [50], to primary human bronchial epithelia [58]), there is evidence that CFTR correction involves biosynthetic pathway and quality control components that are cell-type specific [59]. Immortalized overexpressing cell-lines, even those derived from human bronchial epithelia, do not always predict drug activity in primary cultures for corrector compounds [20]. Thus, especially when addressing questions focusing on biogenesis with potential translational impact, studies using our assay will need to be complemented and confirmed by research using material better recapitulating *in vivo* cellular processing. This has been the approach followed for the currently approved correctors VX-809 and VX-661, modifications of hits first identified using an overexpressing mouse fibroblast cell-line [60].

#### Accurate measurements of low CFTR ion channel function

In addition to membrane proximity, our assay quantifies channel function. Here we confirm previously published data, showing how two different protocols - one measuring the maximal rate of I^−^ entry (*Δ*[*I^−^*]in/*Δt*) during CFTR activation [23], and the other estimating CFTR conductance by fitting quenching time course after steady-state activation is reached [38] – provide results which are consistent with those obtained with other techniques (e.g. Ussing chamber short-circuit current measurements, high-throughput electrophysiology). Thus both *G*_CFTR_ (Figure 4B-D, Supporting Figure S8) [40–42] and (*Δ*[*I^−^*]_in_/*Δt*) (Figures 3 and 5B) [21] can accurately estimate CFTR ion channel function. In this study the assay conditions were not optimized to measure high CFTR activities and some measurements hit the upper limit of its dynamic range (e.g. for WT-CFTR, Figs. 3 and 4, Supporting Table S3). If needed, conditions can be altered to avoid assay saturation (e.g. by using lower concentrations of forskolin or I^−^_out_).

Accurate quantification of low conductance values is advantageous in characterizing drug response by CFTR mutants which have particularly low residual activity. For instance, our assay detects strong VX-770 potentiation for R347P-, N1303K- and H1085R-CFTR (Figure 4D and 5D), genotypes giving no significant potentiation over baseline in a Vertex Pharmaceuticals study to profile VX-770 sensitivity [40]. Our results on N1303K are consistent with patch-clamp and other short-circuit current measurements demonstrating effective potentiation of N1303K-CFTR by VX-770 [61–63]. Despite short-circuit current in FRT cells being increased only to less than the 5% of WT-CFTR threshold [40], caution is required in classifying such mutants as “unresponsive” to VX-770, as they might benefit from therapies combining VX-770 with other modulators [62,63]. Equally promising for possible studies on synergistic modulator effects are L927P- and H1045D-CFTR channels, which, because of very low baseline levels give potentiated short-circuit currents only slightly above the 5% of WT-CFTR threshold [40], but are also powerfully potentiated (Figure 4D and 5D).

### Considerations on VX-770 mechanism of action

Our empirical profiling of the VX-770 response in the rare-mutation panel can generate hypotheses on mechanism of action. Considering the sites of mutations resulting in the highest efficacy (fold-potentiation >20, Figure 5D), these appear to link the ATP molecule bound at site 1 (comprising Walker motifs of NBD1, and signature sequence of NBD2) to regions close to the narrowest portion of the permeation pathway, thought to constitute the CFTR gate [64,65], and positioned adjacent to the recently identified VX-770 binding site [17] (Figure 7).

**Figure 7.**
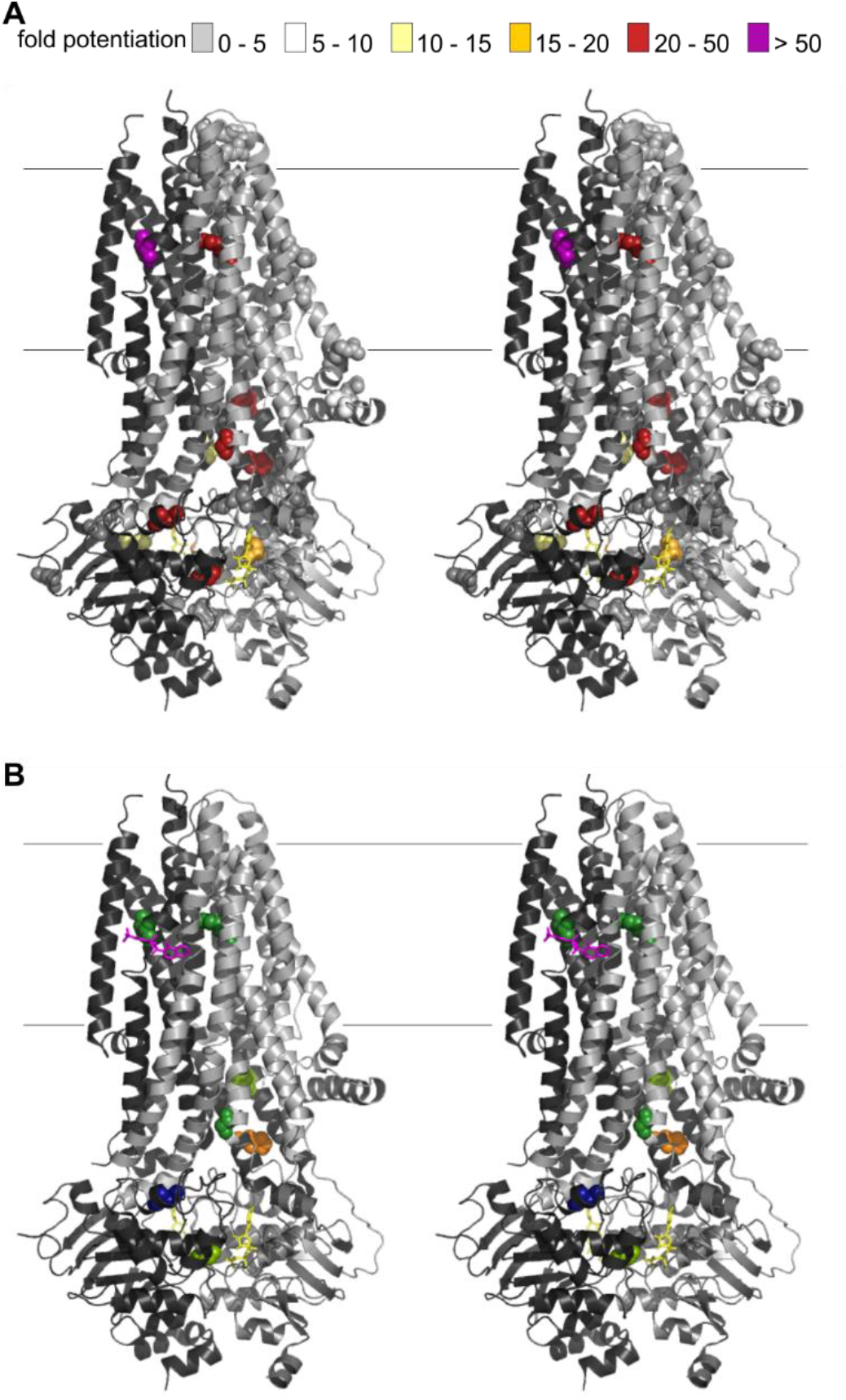
Mapping VX-770 sensitivity on cryo-EM structures. (**A**) Cartoon representation (cross-eye stereo) of phosphorylated, ATP-bound human CFTR (6MSM [18]), with atoms of missense mutations included in the panel highlighted as spheres. Colours indicate degree of fold-potentiation by VX-770. TMD1-NBD1 in light grey; TMD2-NBD2 in dark grey. Fine horizontal lines show approximate position of membrane boundary. (**B**) Only missense mutation sites with most efficacious VX-770 potentiation are shown. Magenta sticks show position of bound VX-770 in 6O2P structure [17]. In cartoon representation, 6O2P and 6MSM are virtually identical (RMSD 0.14 Å, [17]) Mutation-site residues are colour-coded as in Fig. 4 (moving from cytosol to extracellular): G1349, light green; N1303, dark blue; H1054, orange; G178, forest; H1085, light green; R347, forest right; L927 forest left.

Among the highly VX-770-sensitive mutations, all those at the NBD1/NBD2 and NBD/TMD interfaces - introduce charged side chains which would interact unfavourably with other close charges in the conformation observed for phosphorylated, ATP-bound human CFTR (carrying the open-state stabilizing E1371Q mutation, 6MSM, [18] – see Supporting Table S10). Mutations at these sites will particularly destabilize conformations with tight NBD1/NBD2 and NBD/TMD interfaces, such as the NBD-dimerized, open channel conformation [66]. A destabilization of this ABC-canonical open channel conformation is thus likely the cause of the low conductance measured after baseline activation in these mutants. Consistent with this interpretation, N1303K-CFTR channels appear to have almost completely lost the coupling between NBDs and TMDs that normally controls gating, and the rare openings observed are not linked to ATPase cycles at the NBDs [63]. The fact that for all these mutants conductance is greatly increased by VX-770 suggests that drug binding might allow channels to reach an alternative, particularly stable, open state, with a different conformation at the interfaces surrounding site 1.

It has been recently suggested that “undocking” of NBD1 from the TMDs might occur physiologically [67] and several published observations are consistent with the non-canonical VX-770-bound open state described above comprising an undocked NBD1. First, binding of a drug on the MsbA bacterial ABC exporter, at a site not far from the transmembrane VX-770 binding site on CFTR, leads to a distortion of transmembrane helix 4 (TM4) which results in a displacement of the NBD, undocking it from the conserved network of molecular interactions generally stabilizing the NBD/TMD interface [68]. Second, the coupling helix in intracellular loop 4 (ICL4), buried in the NBD1 socket in all the CFTR structures so far reported, was identified as the region for which VX-770 binding decreases hydrogen/deuterium exchange most [69], as would be expected if this helix forms part of a drug-binding site. While the recent cryo-EM structure of the E1371Q-CFTR mutant in complex with VX-770 shows density for only one VX-770 molecule, bound in the transmembrane region [17], it is possible that the exceptionally stable ABC-canonical open conformation of this Walker B mutant [66] prevents NBD1 undocking and thus VX-770 access to a second binding site at the ICL4/NBD1 interface. A second VX-770 binding site, exposed by infrequent undocking of NBD1, would also be consistent with the very slow activation time-course observed upon addition of VX-770 to WT-CFTR, likely reflecting VX-770 having to sequentially occupy two sites before the *P*_O_ can increase [50]. Third, millimolar concentrations of ATP prolong open dwell-times in the presence, but not absence, of VX-770 [70]. This suggests that ATP can bind to/dissociate from a low affinity binding site present on the VX-770-bound open channel conformation. While the ATP binding sites are occluded in the ABC-canonical open channels [71,72], VX-770 induced NBD1 undocking would be expected to alter the NBD interface, possibly resulting in solvent exposure of one of the ATP-binding sites.

The remaining two highly VX-770-sensitive mutations we identify are not at domain interfaces, but close to the CFTR gate: R347P and L927P (Figure 5D, Figure 7). Both mutations replace native sidechains with prolines, which restrict backbone flexibility [73]. R347, in TM6, is important for maintaining a stable conducting pathway [74–76], while L927 is in the unwound segment of TM8 [77,78], underlying CFTR’s unique channel function [78]. The very low conductance measured after baseline activation in both these mutants, suggests that backbone flexibility at both these sites is required for normal channel opening and/or to maintain an open permeation pathway [18]. VX-770 has been hypothesized to increase conformational flexibility of CFTR overall [29]. It is possible that the VX-770 molecule bound at the lipid-CFTR interface might locally (A928 is part of the VX-770 binding site [17]) increase flexibility, facilitating rearrangement of the helices and allowing adoption of the alternative open state described above.

### Implications for pharmacological research

The main advantage of our assay consists in providing simultaneous measurements of ion channel function and biogenesis. Being able to monitor how compounds or mutations affect both number of channels at the membrane and conductance can allow deconvolution of effects on processing from those influencing gating and permeation of the channel. Describing each CF-causing mutation with two coordinates (*ρ* and *G*_CFTR_) is a more informative way of characterizing mutations (e.g. Figure 6) and how they respond to drugs (e.g. Figure 5D), than using either functional or surface-exposure measures alone. The higher information content of measurements will accelerate discovery in projects investigating molecular mechanisms. For instance, using mutagenesis to scan secondary structure elements or to target residues in putative drug-binding sites, hypotheses can be generated or tested rapidly, and results will pinpoint areas worthy of further investigation by more labour-intensive techniques (e.g. patch-clamp/molecular dynamics).

In addition to providing a valuable tool for basic science investigation, our assay could also have a translational impact. While other functional assays, in more native systems (e.g. short-circuit current measurements on primary human bronchial epithelia, forskolin induced swelling of intestinal organoids [79]), will remain fundamental for pre-clinical testing of CFTR-targeting drugs, our assay can usefully complement these.

First, the assay could be useful for development of better precision medicines for CF treatment. Each of the CFTR variants associated with CF could idiosyncratically affect folding, trafficking, stability, gating dynamics and/or permeation - as well as how these properties respond to modulator drugs. A number of modulators are currently approved or in the development pipeline, and therapies combining multiple correctors and potentiators appear to be most effective, at least for patients carrying the F508del mutation [15,16,80]. However, potentiators can negatively interfere with corrector action, and drug-drug interactions are genotype specific [29,30,49]. Because each mutation, other than F508del, is extremely rare, pre-clinical studies using our assay could provide a first molecular characterization of how individual CFTR variants respond to modulator drugs, and drug combinations, in controlled, simplified conditions. Such data can be very valuable to inform drug development, trial design, and therapy choice, especially for genotypes found only extremely rarely in the population [81].

Second, the assay could help develop very effective dual-activity modulator drugs for CF treatment. Both gating/permeation and processing defects likely stem from impaired folding, at least for the common F508del-CFTR variant [82]. However, practical implementation of distinct potentiator and corrector screens might have so far biased the drug development process by selecting compounds for improvement only in one dimension [83]. Screening using our integrated assay, by maintaining the requirement for simultaneous reduction of both defects, will maximise the chances of identifying ligands capable of redressing the primary folding defect. By shifting therapy closer to the root cause of disease, such a drug would likely reduce the need for prevention/treatment of comorbidities and exacerbations, as well as decrease the likelihood of long-term safety and tolerability problems.

Finally, CFTR plays an important role controlling fluid movement across several epithelia [2,84], and it has been implicated in a number of pathologies, including secretory diarrhoeas [85], COPD [86,87], polycystic kidney disease [88] and others [89,90]. It is likely that, given the complexity of CFTR folding [8,82], many CFTR-targeting compounds will alter its cellular processing (e.g. [10]), suggesting that the assay could also be usefully deployed as part of the development of novel CFTR-targeting compounds for treatment of other diseases, beyond CF.

### Experimental Procedures

#### Construction of the pIRES2-mCherry-YFPCFTR plasmid

The pIRES2-mCherry-YFPCFTR plasmid was obtained with two sequential subcloning steps. First, a 1.727 kb region of pcDNA3.1-YFP-CFTR [23], containing the YFP-coding sequence, was subcloned into pIRES-eGFP-CFTR, a gift from David Gadsby (Rockefeller University), using the NheI and BlpI restriction sites. Subsequently a 0.737 kb region from pIRES2-mCherry-p53 deltaN ([91], Addgene), containing the mCherry-coding segment and part of the IRES, was subcloned into the pIRES-eGFP-YFPCFTR plasmid using the NotI and BmgBI/BtrI restriction sites. This resulted in the pIRES2-mCherry-YFPCFTR plasmid, with the IRES2 positioned between the two open reading frames for YFP-CFTR and mCherry.

To generate the rare-mutation panel, point mutations were introduced in the pIRES2-mCherry-YFPCFTR plasmid using site-directed mutagenesis (Quikchange protocol, Stratagene).

#### HEK293 cell culture, transfection and incubation

HEK293 cells were maintained in Dulbecco’s modified Eagle’s medium (DMEM), supplemented with 2 mM L-glutamine, 100 U/ml penicillin and streptomycin, and 10% fetal bovine serum (all Life Technologies). Cells were seeded in poly-D-lysine-coated, black-walled 96-well plates (Costar, Fisher Scientific), and transiently transfected with the pIRES2-mCherry-YFPCFTR plasmid using Lipofectamine 2000 (Life Technologies), following manufacturer instructions. After transfection, cell plates were returned to the 37 °C incubator for 24 hours. Prior to imaging, plates were incubated for another 24 hours, at 37 °C or 28 °C, in 100 μl DMEM including DMSO (vehicle), 10 μM VX-809, or 10 μM VX-770 plus 10 μM VX-809 (Selleck Chemicals), as indicated. The assay is currently run using 96-well plates but small changes could make it compatible to a 384 well plate format.

#### Image acquisition

Before imaging, cells were washed twice with 100 μl standard buffer (140 mM NaCl, 4.7 mM KCl, 1.2 mM MgCl_2_, 5 mM HEPES, 2.5 mM CaCl_2_,1mM glucose, pH 7.4). The ImageXpress Micro XLS (Molecular Devices), an automated inverted wide-field fluorescence microscope with a temperature-controlled chamber (set to 37 °C or 28 °C, as indicated), was used for image acquisition. Protocols for automated fluid additions, enabled by a robotic arm, were created using MetaXpress software (Molecular Devices). For imaging of YFP-CFTR, a 472 ± 30 nm excitation filter, and a 520 ± 35 nm emission filter were used. Excitation/emission filters at 531 ± 20 nm and 592 ± 20 nm were used for imaging of mCherry.

For localization of CFTR, a 60× objective was used to take 9 16-bit images per well of both fluorophores. To evaluate CFTR function, a 20× objective was used. Two 16-bit images of mCherry were taken, one at the start and one the end of the protocol. In addition, 16-bit images of the YFP fluorescence, were taken at an acquisition frequency of 0.5 Hz. For the I^−^ first protocol ((A), see below), after 20 s, 50 μl of 300 mM I^−^ buffer (300 mM NaI, 4.7 mM KCl, 1.2 mM MgCl_2_, 5 mM HEPES, 2.5 mM CaCl_2_,1mM glucose, pH 7.4) was added to the standard buffer, so that the final concentration of I^−^ in the extracellular medium was 100 mM. Another 40 s later, a further 50 μl of a 100 mM I^−^ buffer containing 40 μM forskolin (100 mM NaI, 4.7 mM KCl, 1.2 mM MgCl_2_, 5 mM HEPES, 2.5 mM CaCl_2_,1mM glucose, 40 μM forskolin, pH 7.4) was added, so that the final concentration of forskolin in the extracellular medium was 10 μM, while concentration of other components remained unaltered. For the I^−^ last protocol ((B), below), after 20 s of imaging, CFTR was activated, in the absence of extracellular I^−^, by addition of 50 μl standard buffer containing activating compounds (forskolin or forskolin + VX-770 both to reach final concentrations of 10 μM). After a further 230 s, by which time CFTR is assumed to be gating at steady state [38], extracellular I^−^ was raised to 100 mM (final concentration) by adding 50 μl of I^−^ buffer (as standard buffer with 140 mM NaCl replaced with 400 mM NaI). Images were taken for another 40 s. Activating compounds were also included in the second addition so as not to alter final extracellular concentrations.

#### Image analysis

Image analysis was automated using MATLAB mathematical computing software (MathWorks). Separate analysis protocols were implemented to estimate CFTR membrane proximity and ion channel function.

##### CFTR membrane proximity

First, mCherry images were binarized, and basic morphological operations (opening, closing, area opening, and dilation) were carried out to reduce noise. A distance transform with locally imposed minima was used to segment images by means of a watershed transformation, and define cell boundaries. Cells were removed from analysis if they had an area of under 108 μm^2^ or over 5400 μm^2^, if they had a major axis length of less than 32.4 μm, if the area over perimeter was less than 25 or over 300, and if they were touching the edge of the image. A 1.08 μm band, 10 or 5 pixels wide (depending on the resolution of the image), within the border of each cell was defined as the membrane-proximal zone.

Background was selected by inverting the binarized and morphologically opened mCherry image, after which it was morphologically closed using a large structuring element, to prevent cells from being selected as background. Average background intensity was then subtracted from each pixel, and the YFP and mCherry fluorescence intensity of each cell was normalized to the median YFP and mCherry fluorescence intensities of cells expressing WT-CFTR on the same plate. If the average normalized fluorescence intensity fell below 0 (due to low transfection efficiency and high background noise), cells were removed from analysis.

In order to estimate CFTR membrane proximity for each cell (defined as *ρ*, see Results), the average normalized YFP fluorescence intensity within the membrane-proximal zone was divided by the average normalized mCherry fluorescence over the entire cell.

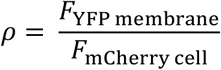

##### CFTR ion channel function

For assessment of CFTR function, two different protocols were used. For both, cells were selected based on the mCherry fluorescence images that were taken at the beginning and at the end of the protocol. The images were binarized using an adaptive threshold, after which they were dilated and combined to account for possible minor movement of cells during the time course.

###### (A) I^−^ first Protocol

The fluorescence at the time point before addition of I^−^ was used to normalize YFP fluorescence intensity. The concentration of I^−^ inside the cells ([*I*^−^]_in_) can be estimated with the following equation [23], in which the binding affinity for I^−^ (*K*_I_) to YFP(H148Q/I152L) is set to 1.9 mM [22] and the normalized fluorescence intensity over time (*F*(*t*)) is determined experimentally.

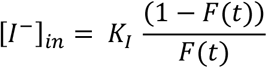

Data is collected every 2 seconds, so the change [*I*^−^]_in_ observed at each time point can be estimated and used to calculate the rate of I^−^ entry (in mM/s):

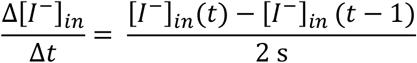

The maximal observed rate of I^−^ entry is used as a measure of cellular anion conductance. To determine whether there was increased CFTR-mediated anion conductance, the maximal rate of I^−^ entry after addition of forskolin (which activates CFTR due to increased phosphorylation by cAMP-dependent protein kinase), was compared to the maximal rate of I^−^ entry after addition of DMSO (vehicle, negative control).

###### (B) I^−^ last Protocol

CFTR activation (by addition of 10 μM forskolin with or without 10 μM VX-770, as indicated) was first allowed to reach steady state in the absence of I^−^ and quenching of YFP in the 40 s following extracellular I^−^ addition was measured. A simple mathematical model was used to fit observed fluorescence quenching, and estimate CFTR conductance as described [38]. Briefly, the model includes four free parameters: CFTR conductance at steady-state (*G*_CFTR_), membrane potential at steady-state, immediately prior to I^−^ addition (*V*_M_), and conductance (*G*_trans_) and time constant (τ_trans_) of a transient, endogenous non-CFTR anion conductance. The values of the four parameters were estimated by minimizing the sum of squared residuals obtained by comparing the time course of the observed average fluorescence intensity within cells to the proportion of anion-free YFP chromophore predicted by the model (both normalized to the time point before I^−^ addition). However, when the quenching time course was too fast and did not provide enough information to uniquely identify all four parameters, the value of the latter two parameters (*G*_trans_ and τ_trans_) was constrained to the average values obtained with negative controls, and only *G*_CFTR_ and *V*_M_ were left free to vary [38]. Experimental data are well described by the model, suggesting that YFP chromophore molecules, whether fused to CFTR inserted in intracellular vesicles or in the plasma membrane, behave as a single population.

For both protocol (A) and (B) the value obtained from analysis of the observed YFP-CFTR fluorescence quenching (*G*_CFTR_ and (*Δ*[*I*^−^]_in_/*Δt*) respectively) was corrected to account for variations in transfection efficiency. Thus, the metric reporting ion channel function was normalised for each condition/genotype by dividing by the mean *F*_mCherry_ within the cell selection (which, in turn, was normalized to *F*_mCherry_ measured for WT in the same plate).

#### Statistical analysis

To determine whether the observed differences in *ρ*, (*Δ*[*I*^−^]_in_/*Δt*) or *G*_CFTR_ resulting from experimental manipulation and/or mutations were statistically significant, we performed either independent or paired t-tests (pairing different genotypes/conditions measured in the same multi-well plate). When required, either a Bonferroni or a Benjamini-Hochberg correction was applied to adjust for multiple comparisons. Data in graphs represent mean ± SEM, and the significance level was pre-specified as α = 0.05. Statistical analysis was carried out using MATLAB (MathWorks), SigmaPlot (Systat Software), SPSS (IBM), or Excel (Microsoft).

#### Data availability statement

Most data is presented in the main-article Figures. In addition, the Supporting Information includes: information on the statistical analyses performed (Tables S1-S4, S6, S7); paired t-tests plots and distributions of log_10_*ρ* values for each mutant in the rare-mutation panel (Figure S5); a comparison between our results for the rare-mutation panel and published data (Figure S8).

Analysis code and example images to run it on are provided for readers. All the necessary instructions and files can be found at: https://github.com/stellaprins/CFTRimg

## Non-standard Abbreviations

ABC: ATP-binding cassette
CF: Cystic Fibrosis
CFTR: Cystic Fibrosis Transmembrane Conductance Regulator
FRT: Fischer Rat Thyroid
*F*_mCherry cell_: average normalized mCherry fluorescence intensity over the entire cell
*F*_YFP cell_: average normalized YFP fluorescence intensity over the entire cell
*F*_YFP membrane_: average normalized YFP fluorescence intensity within the membrane-proximal zone
*G*_CFTR_: CFTR conductance
*G*_trans_: transient anion conductance
IRES: internal ribosome entry site
NBD: nucleotide binding domain
*P*_O_: open probability
*ρ*: CFTR membrane proximity, as defined in this paper
SSR: sum of squared residuals
τ_trans_: time constant of the transient anion conductance
TM: transmembrane helix
TMD: transmembrane domain
*V*_M_: membrane potential, after steady state activation of CFTR
WT: wild type
YFP: yellow fluorescent protein

## Acknowledgements

We thank Dr William Andrews, Central Molecular Laboratory, UCL for help with molecular biology. We are also grateful to Sam Ranasinghe and staff at the UCL Confocal Imaging Facility, Division of Biosciences, for their help with the temperamental equipment.

## Funding and additional information

EL was supported by grant 15UCL04, funded by the Sparks charity and Cystic Fibrosis Trust. SP was supported by grant SRC005 funded by the Cystic Fibrosis Trust. CH was supported by EPSRC grant EP/F500351/1, and ACS was awarded a British Pharmacological Society Vacation Studentship.

