## Supplementary for "Fluorescence assay for simultaneous quantification of CFTR ion-channel function and plasma membrane proximity"

### Supporting Information

**Supporting Table S1** Paired Sample t-tests comparing the $\log_{10}\rho$ of F508del-CFTR or F508del/R1070W-CFTR after different chronic incubation conditions. An independent t-test was performed to assess the significance of the difference in $\log_{10}\rho$ of WT-CFTR and F508del-CFTR at 37 °C and 28 °C. P-values were Bonferroni adjusted to account for multiple comparisons.

| | | Mean | SD | SD for mean difference | df | T value | Adjusted P value |
|---|---|---|---|---|---|---|---|
| 37°C | F508del | -0.59 | 0.05 | 0.02 | 6 | -4.64 | 0.01 |
| | F508del + VX-809 | -0.55 | 0.06 | | | | |
| | F508del + VX-809 | -0.59 | 0.07 | 0.02 | 5 | 4.22 | 0.02 |
| | F508del + VX-809 + VX-770 (c) | -0.63 | 0.06 | | | | |
| | F508del | -0.58 | 0.07 | 0.02 | 7 | -5.78 | 2.01E-03 |
| | F508del/R1070W | -0.53 | 0.08 | | | | |
| 28°C | F508del | -0.51 | 0.09 | 0.04 | 6 | -12.23 | 6.00E-05 |
| | F508del + VX-809 | -0.31 | 0.07 | | | | |
| | F508del + VX-809 | -0.34 | 0.08 | 0.20 | 10 | 4.06 | 4.54E-03 |
| | F508del + VX-809 + VX-770 (c) | -0.50 | 0.09 | | | | |
| | F508del | -0.54 | 0.07 | 0.13 | 6 | -4.73 | 0.01 |
| | F508del/R1070W | -0.40 | 0.10 | | | | |
| temperature correction | 37°C (WT - F508del) | 0.54 | 0.08 | NA | 44 | 5.59 | 4.05E-06 |
| | 28°C (WT - F508del) | 0.41 | 0.08 | | | | |

**Supporting Table S2** Independent t-tests comparing the maximal rate of $I^-$ entry after addition of 10µM forskolin vs. after DMSO (control). Cells were transfected with WT-CFTR, F508del-CFTR, or F508del/R1070W-CFTR and incubated at either 37 °C or 28 °C, with or without 10 µM VX-809, 24 hours before imaging. In some conditions the potentiator VX-770 (10 µM) was added acutely together with forskolin.

| | | VX-809 | VX-770 (a) | | Mean | SD | df | T value | P value |
|---|---|---|---|---|---|---|---|---|---|
| WT | 37°C | - | - | Forskolin | 0.180 | 0.059 | 16 | 8.65 | 1.99E-07 |
| | | | | DMSO | 0.010 | 0.003 | | | |
| | | - | + | Forskolin | 0.214 | 0.056 | 4 | 6.36 | 3.13E-03 |
| | | | | DMSO | 0.008 | 0.003 | | | |
| | 28°C | - | - | Forskolin | 0.137 | 0.023 | 10 | 13.18 | 1.20E-07 |
| | | | | DMSO | 0.010 | 0.005 | | | |
| | | - | + | Forskolin | 0.214 | 0.056 | 5 | 6.36 | 3.13E-03 |
| | | | | DMSO | 0.008 | 0.003 | | | |
| F508del | 37°C | - | - | Forskolin | 0.009 | 0.005 | 15 | 0.71 | 0.49 |
| | | | | DMSO | 0.007 | 0.004 | | | |
| | | - | + | Forskolin | 0.014 | 0.007 | 4 | 1.50 | 0.21 |
| | | | | DMSO | 0.007 | 0.002 | | | |
| | | + | + | Forskolin | 0.034 | 0.008 | 9 | 5.72 | 2.87E-04 |
| | | | | DMSO | 0.013 | 0.004 | | | |
| | 28°C | - | - | Forskolin | 0.012 | 0.009 | 10 | 0.00 | 1.00 |
| | | | | DMSO | 0.012 | 0.014 | | | |
| | | - | + | Forskolin | 0.059 | 0.042 | 5 | 2.71 | 0.04 |
| | | | | DMSO | 0.004 | 0.001 | | | |
| | | + | + | Forskolin | 0.124 | 0.039 | 9 | 6.67 | 9.12E-05 |
| | | | | DMSO | 0.007 | 0.004 | | | |
| F508del/R1070W | 37°C | - | - | Forskolin | 0.044 | 0.026 | 12 | 2.87 | 0.01 |
| | | | | DMSO | 0.014 | 0.010 | | | |
| | 28°C | - | - | Forskolin | 0.090 | 0.032 | 6 | 4.38 | 4.68E-03 |
| | | | | DMSO | 0.015 | 0.014 | | | |

**Supporting Table S3** Independent t-tests comparing the maximal rate of $I^-$ entry after addition of 10 µM forskolin in varying conditions. P-values were Bonferroni adjusted to account for multiple comparisons.

| | Mean | SD | df | T value | P value | adjusted P value |
|---|---|---|---|---|---|---|
| WT 37°C | 0.180 | 0.059 | 10 | 0.88 | 0.40 | 0.80 |
| WT + VX-770 (a) 37°C | 0.214 | 0.056 | | | | |
| WT 28°C | 0.137 | 0.023 | 8 | 2.71 | 0.03 | 0.05 |
| WT + VX-770 (a) 28°C | 0.207 | 0.058 | | | | |
| WT 37°C | 0.180 | 0.059 | 13 | 1.70 | 0.11 | 0.22 |
| WT 28°C | 0.137 | 0.023 | | | | |
| WT + VX-770 (a) 37°C | 0.214 | 0.056 | 5 | 0.17 | 0.87 | 1.00 |
| WT + VX-770 (a) 28°C | 0.207 | 0.058 | | | | |
| F508del + VX-770 (a) 37°C | 0.014 | 0.007 | 6 | 3.52 | 0.01 | 0.02 |
| F508del + VX-770 (a) + VX809 37°C | 0.034 | 0.008 | | | | |
| F508del + VX-770 (a) 28°C | 0.059 | 0.042 | 7 | 2.34 | 0.05 | 0.10 |
| F508del + VX-770 (a) + VX809 28°C | 0.124 | 0.039 | | | | |
| F508del + VX-770 (a) + VX809 37°C | 0.034 | 0.008 | 9 | 5.06 | 6.77E-04 | 1.35E-03 |
| F508del + VX-770 (a) + VX809 28°C | 0.124 | 0.039 | | | | |
| F508del 37°C | 0.009 | 0.005 | 13 | -3.70 | 2.69E-03 | 0.01 |
| F508del/R1070W 37°C | 0.044 | 0.026 | | | | |
| F508del/R1070W 37°C | 0.044 | 0.026 | 9 | -2.62 | 0.03 | 0.06 |
| F508del/R1070W 28°C | 0.090 | 0.032 | | | | |

**Supporting Table S4** Summary of statistical data for membrane density profiling of rare mutation panel.

| Mutation | Mean $\rho$ | SEM | n |
|---|---|---|---|
| **A455E** | 0.327 | 0.027 | 12 |
| **A46D** | 0.363 | 0.022 | 12 |
| **A559T** | 0.371 | 0.047 | 12 |
| **A561E** | 0.328 | 0.018 | 13 |
| **D110E** | 0.827 | 0.044 | 12 |
| **D110H** | 0.649 | 0.034 | 11 |
| **D1152H** | 0.688 | 0.022 | 12 |
| **D1270N** | 0.782 | 0.043 | 11 |
| **D579G** | 0.363 | 0.024 | 12 |
| **E193K** | 0.65 | 0.021 | 13 |
| **E56K** | 0.338 | 0.016 | 11 |
| **E92K** | 0.254 | 0.014 | 12 |
| **F1052V** | 0.761 | 0.031 | 10 |
| **F1074L** | 0.423 | 0.021 | 11 |
| **F508del** | 0.315 | 0.02 | 14 |
| **G1244E** | 0.857 | 0.037 | 11 |
| **G1349D** | 0.747 | 0.029 | 12 |
| **G178R** | 0.69 | 0.017 | 12 |
| **G551D** | 1.042 | 0.034 | 12 |
| **G551S** | 1.167 | 0.033 | 12 |
| **G85E** | 0.281 | 0.012 | 12 |
| **G970R** | 0.815 | 0.045 | 11 |
| **H1054D** | 0.308 | 0.01 | 10 |
| **H1085R** | 0.307 | 0.017 | 9 |
| **I336K** | 0.398 | 0.015 | 10 |
| **I507del** | 0.305 | 0.021 | 11 |
| **K1060T** | 0.632 | 0.025 | 11 |
| **L1065P** | 0.296 | 0.01 | 12 |
| **L1077P** | 0.294 | 0.014 | 12 |
| **L206W** | 0.309 | 0.024 | 12 |
| **L467P** | 0.333 | 0.014 | 13 |

| Mutation | Mean $\rho$ | SEM | n |
|---|---|---|---|
| **L927P** | 0.406 | 0.022 | 11 |
| **M1101K** | 0.294 | 0.015 | 11 |
| **N1303K** | 0.588 | 0.02 | 11 |
| **P67L** | 0.319 | 0.015 | 11 |
| **R1066C** | 0.306 | 0.012 | 11 |
| **R1066H** | 0.29 | 0.013 | 11 |
| **R1066M** | 0.402 | 0.038 | 12 |
| **R1070Q** | 0.899 | 0.041 | 12 |
| **R1070W** | 0.438 | 0.019 | 10 |
| **R117C** | 0.626 | 0.035 | 12 |
| **R117H** | 0.662 | 0.018 | 12 |
| **R1283M** | 0.543 | 0.027 | 11 |
| **R334W** | 0.985 | 0.041 | 12 |
| **R347H** | 1.058 | 0.035 | 12 |
| **R347P** | 0.377 | 0.013 | 13 |
| **R352Q** | 1.046 | 0.033 | 13 |
| **R560S** | 0.307 | 0.02 | 12 |
| **R560T** | 0.332 | 0.019 | 13 |
| **R74W** | 0.522 | 0.022 | 12 |
| **S1251N** | 0.859 | 0.033 | 11 |
| **S1255P** | 0.804 | 0.029 | 12 |
| **S341P** | 0.487 | 0.021 | 11 |
| **S492F** | 0.316 | 0.013 | 12 |
| **S549N** | 0.835 | 0.021 | 11 |
| **S549R** | 0.432 | 0.029 | 11 |
| **S945L** | 0.358 | 0.018 | 10 |
| **S977F** | 0.749 | 0.038 | 11 |
| **T338I** | 0.681 | 0.024 | 13 |
| **V520F** | 0.336 | 0.02 | 12 |
| **Y569D** | 0.298 | 0.015 | 13 |
| **WT** | 0.809 | 0.019 | 23 |

**Supporting Figure S5**

Probability density distributions for $\log_{10}\rho$ values for each CFTR mutant in the panel (orange), compared to WT-CFTR (blue). Plots on left illustrate measurements obtained for WT and mutant from individual plates, paired for statistical analysis. Asterisks indicate a significant difference ($P < 0.05$) in mean $\log_{10}\rho$ between WT and mutant, following paired t-tests and correction for multiple comparisons using Benjamini-Hochberg procedure.

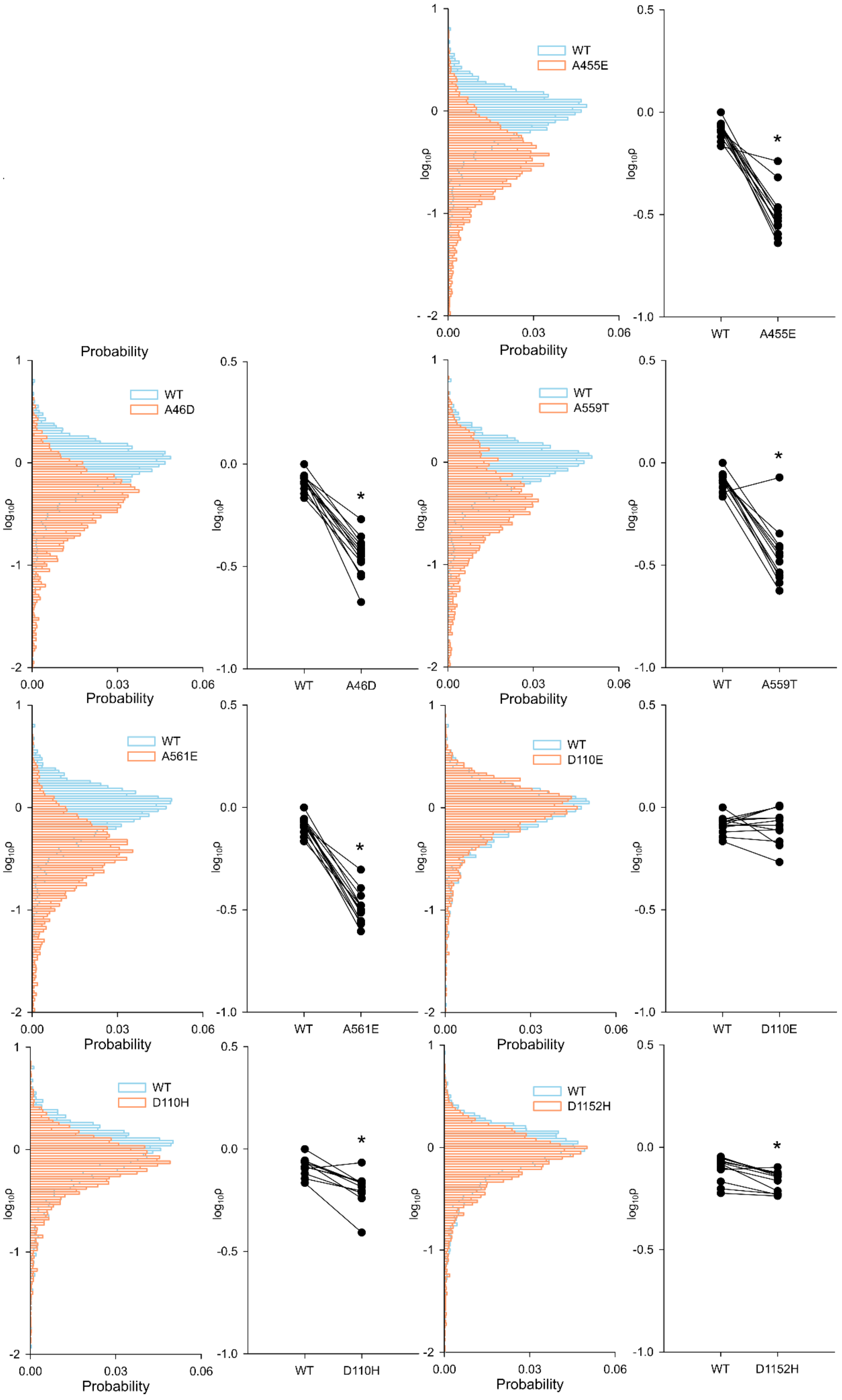
WT
A455E
WT
A46D
WT
A559T
WT
A561E
WT
D110E
WT
D110H
WT
D1152H
Probability
log10ρ
0.00
0.03
0.06
*

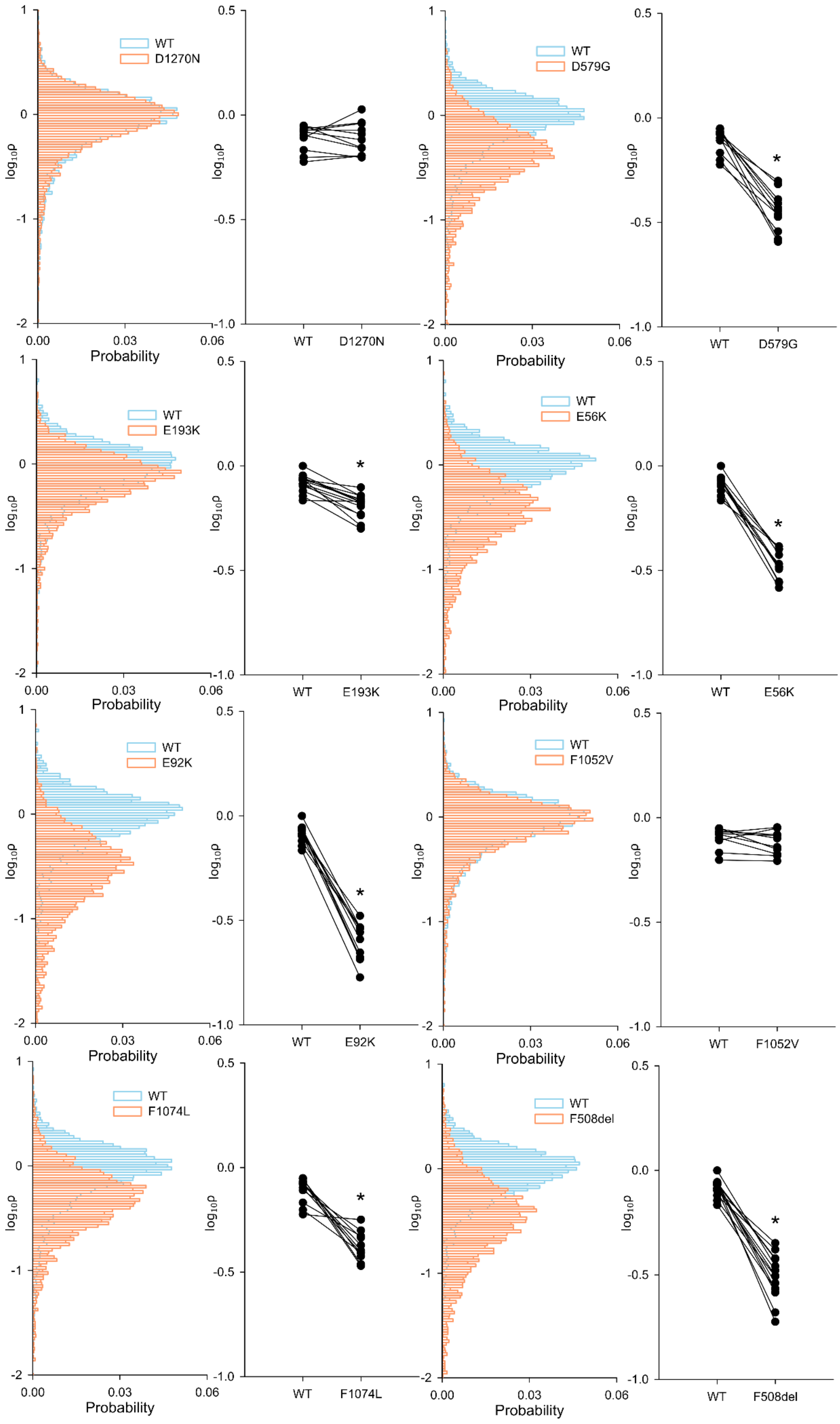
WT
D1270N
D579G
E193K
E56K
E92K
F1052V
F1074L
F508del
Probability
log10ρ
*

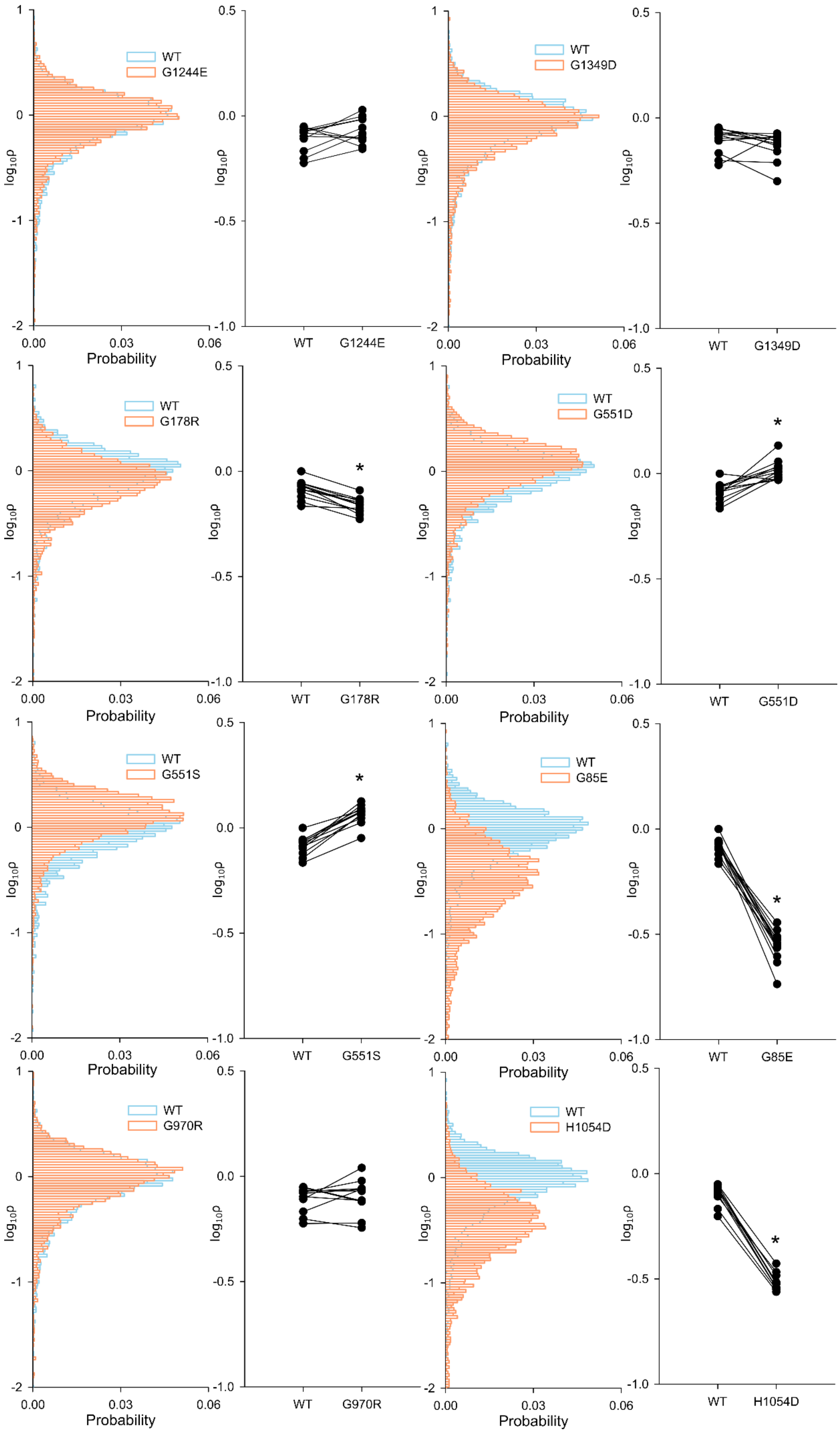
WT
G1244E
G1349D
G178R
G551D
G551S
G85E
G970R
H1054D
log10ρ
Probability
0.00
0.03
0.06
0.5
0.0
-0.5
-1.0
1
0
-1
-2
*

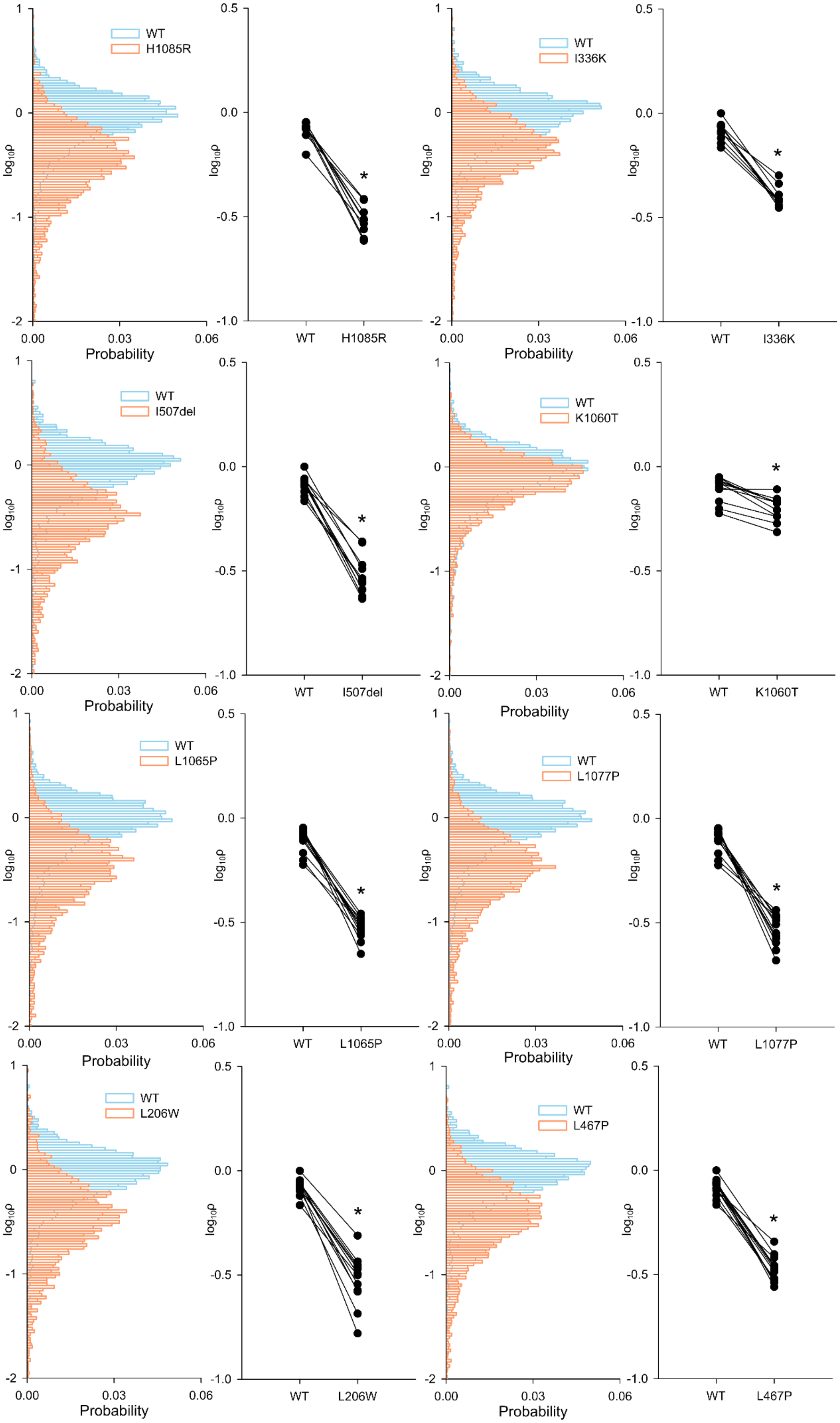
WT
H1085R
I336K
I507del
K1060T
L1065P
L1077P
L206W
L467P
Probability
log10ρ
*

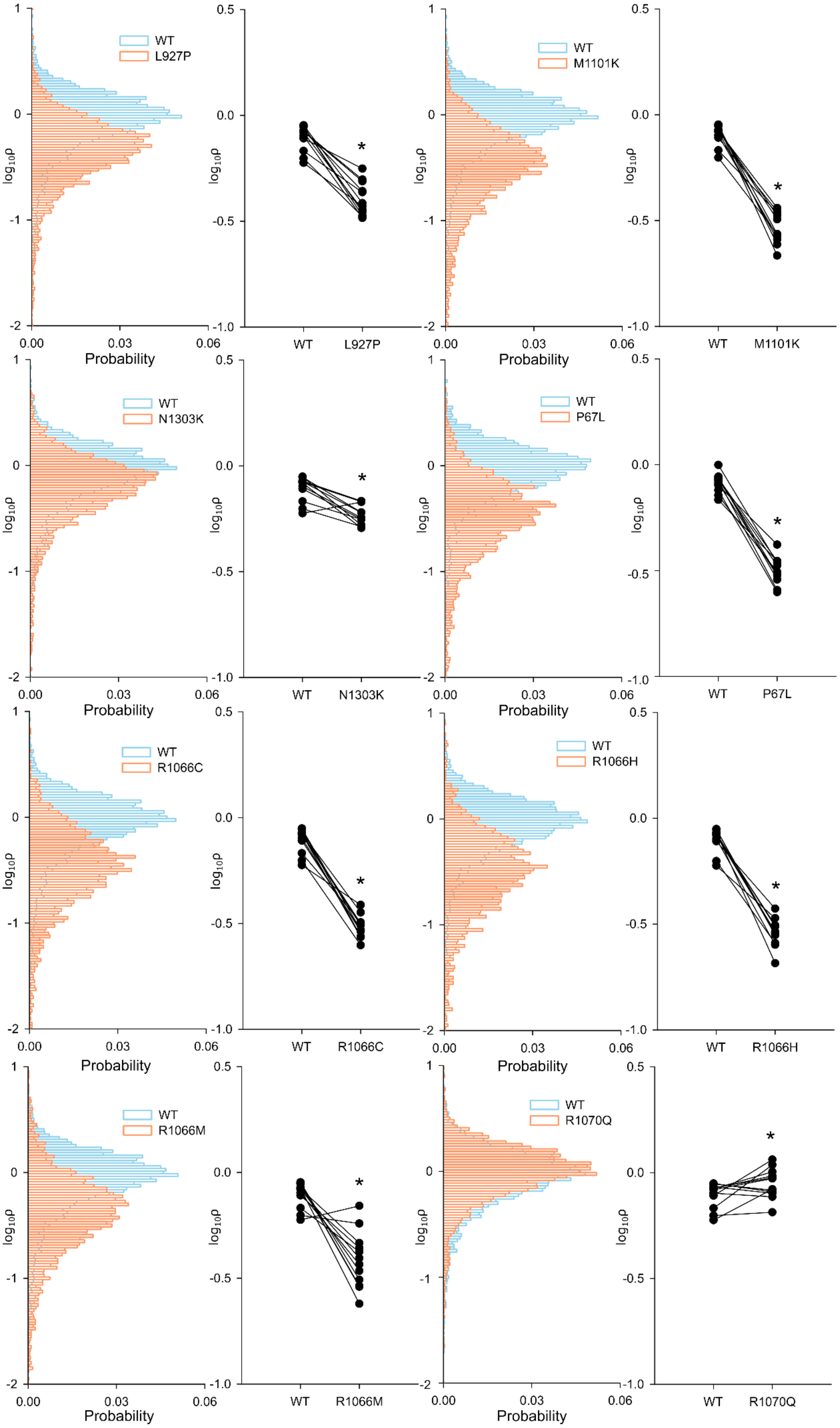
WT
L927P
M1101K
N1303K
P67L
R1066C
R1066H
R1066M
R1070Q
Probability
$\log_{10}\rho$
*

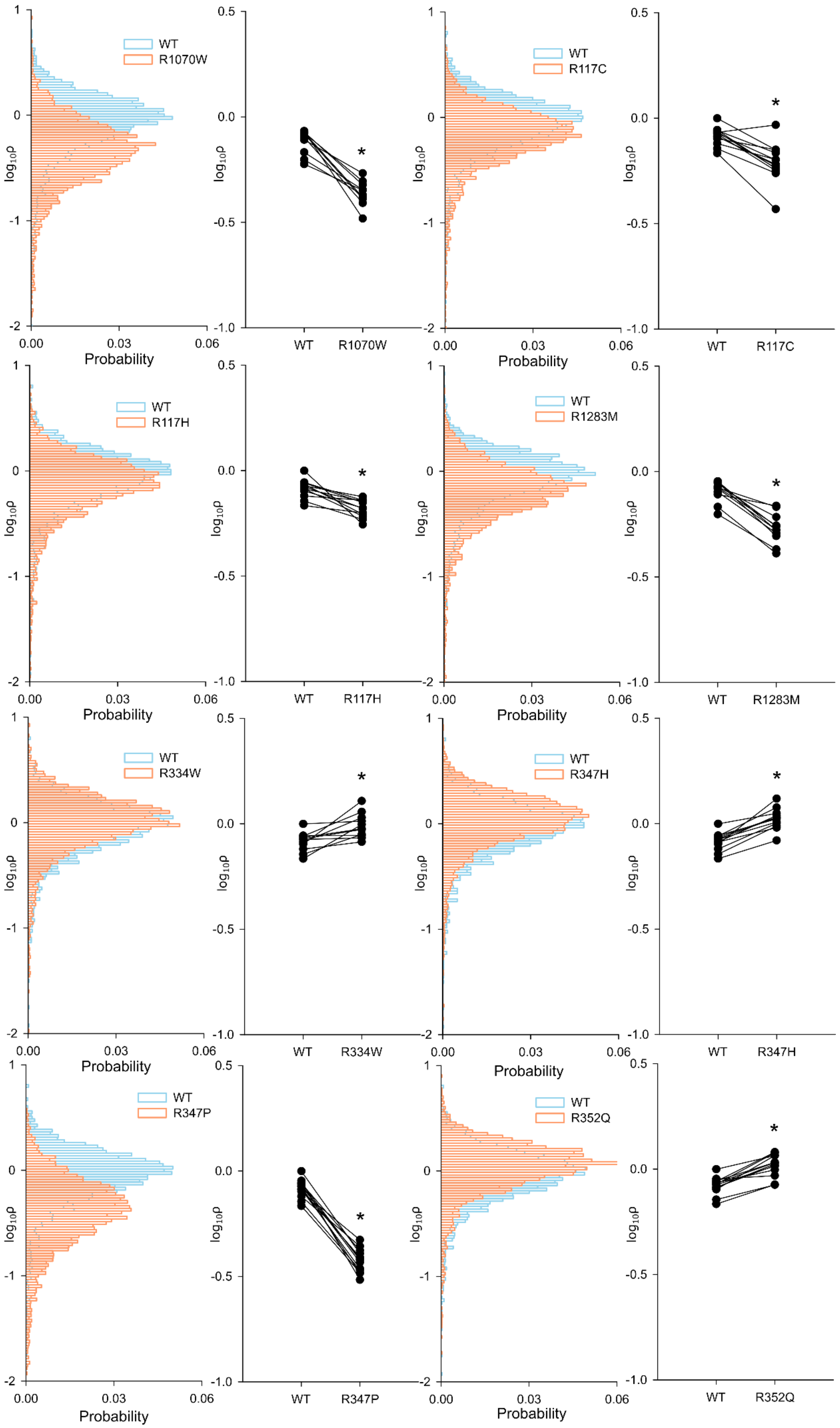
WT
R1070W
log10ρ
Probability
0.00
0.03
0.06
1
0
-1
-2
0.5
0.0
-0.5
-1.0
*
WT
R117C
WT
R117H
WT
R1283M
WT
R334W
WT
R347H
WT
R347P
WT
R352Q

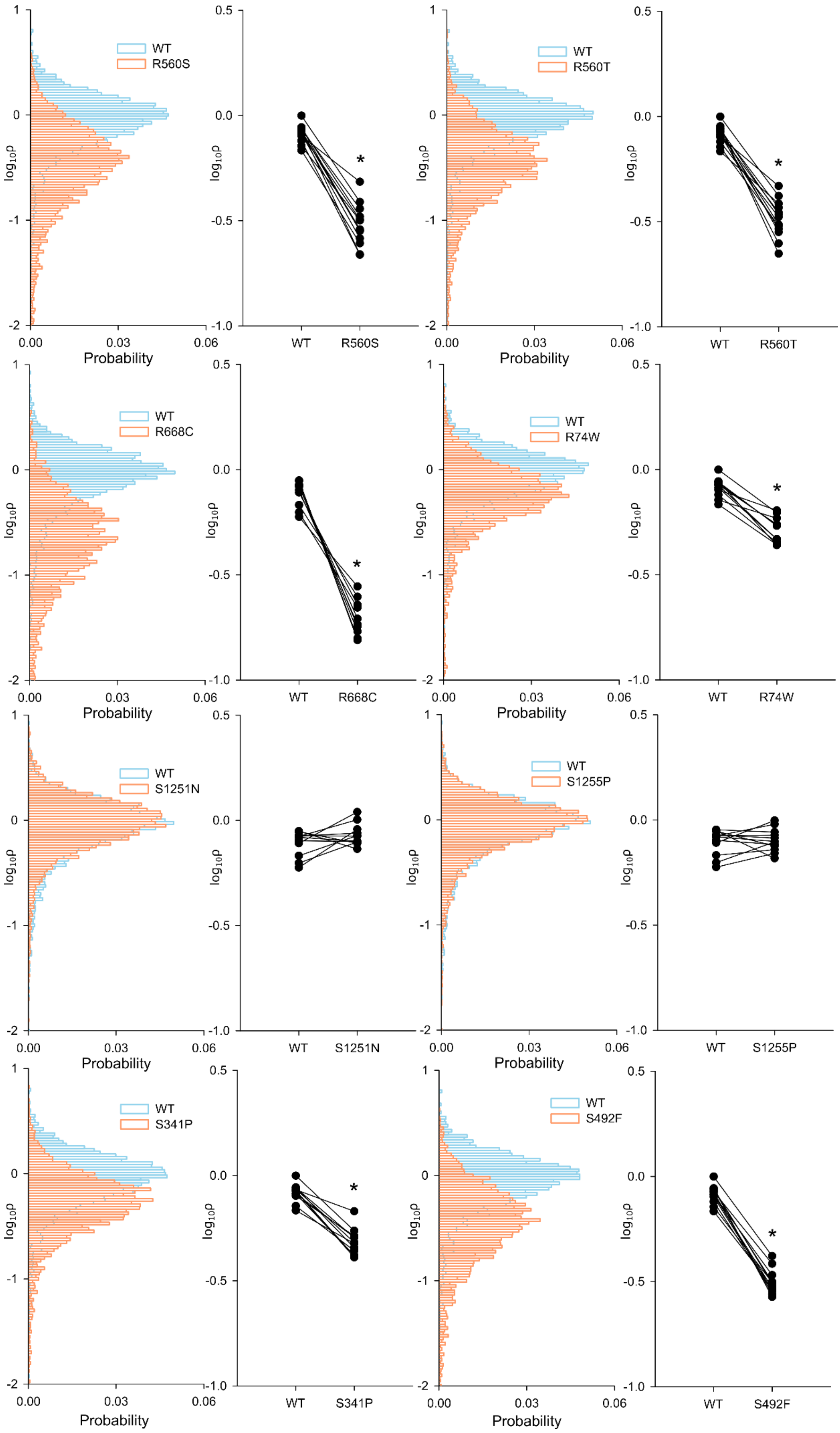
WT
R560S
WT
R560T
WT
R668C
WT
R74W
WT
S1251N
WT
S1255P
WT
S341P
WT
S492F
log₁₀ρ
Probability
*
WT R560S
WT R560T
WT R668C
WT R74W
WT S1251N
WT S1255P
WT S341P
WT S492F

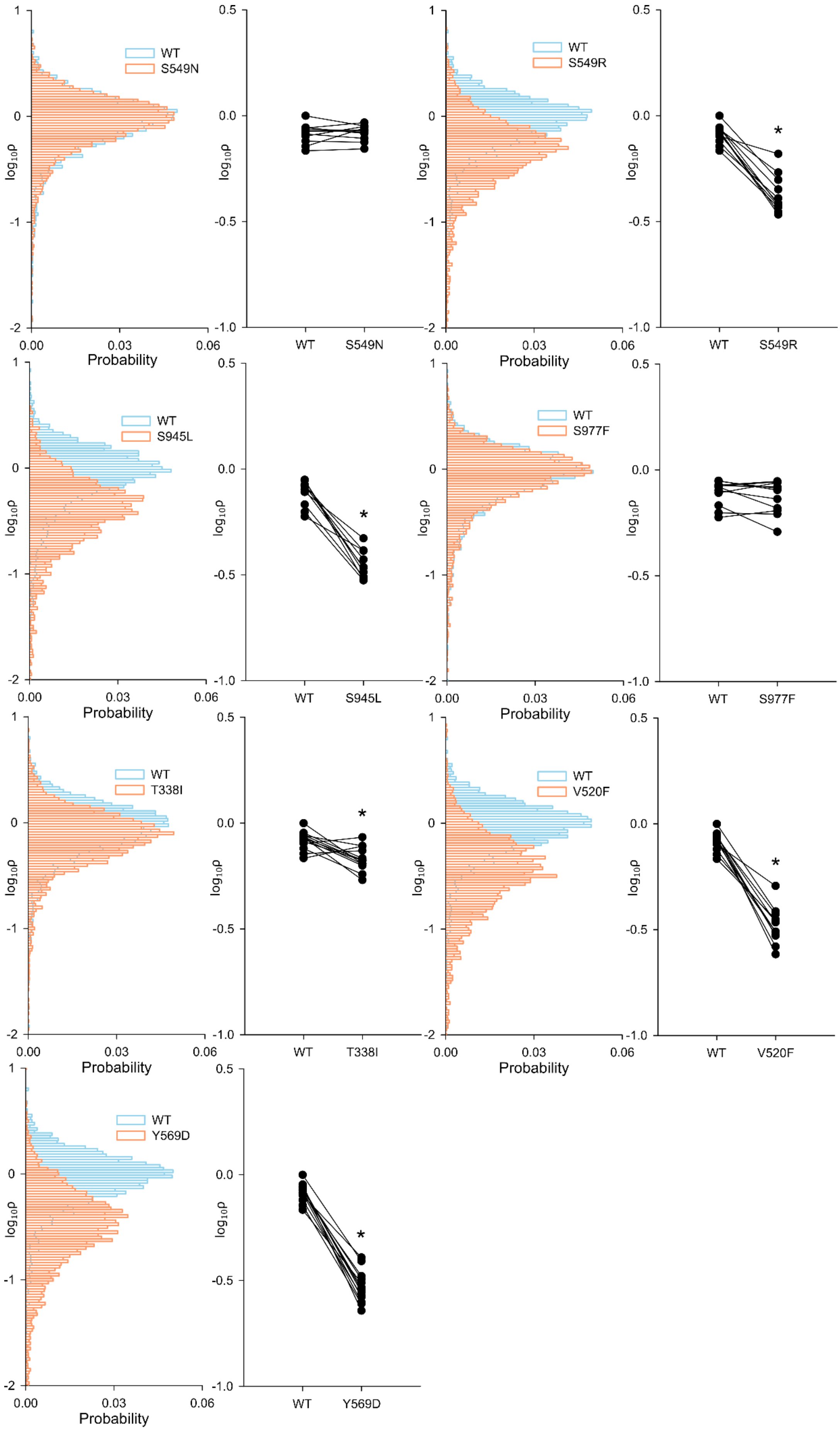
WT
S549N
S549R
S945L
S977F
T338I
V520F
Y569D
log10ρ
Probability
0.00
0.03
0.06
*

**Supporting Table S6**

Data for CFTR conductance ($G_{CFTR}$_norm, nS) profiling of rare mutation panel, in vehicle control conditions with no CFTR activation (**DMSO**), and after baseline activation with 10 μM forskolin (**fsk**). For each genotype, normality (Shapiro-Wilk) and equal variance preliminary tests were performed. Statistical significance of difference between control and baseline CFTR activity was assessed with a one-tailed t-test, assuming equal or unequal (marked with *) variance, depending on result of preliminary test. For R1066H and R1070W, the normality test of input groups failed and significance of within genotype difference between groups was quantified with a non-parametric Mann-Whitney Rank Sum Test.

| Mutation | DMSO | SEM | n | fsk | SEM | n | P value (t test) | P value (Rank Sum test) |
|---|---|---|---|---|---|---|---|---|
| **A455E** | 0.14 | 0.10 | 3 | 0.88 | 0.33 | 3 | 0.0490 | |
| **A46D** | 0.36 | 0.19 | 3 | 2.14 | 1.16 | 3 | 0.1035 | |
| **A559T** | 0.22 | 0.11 | 3 | 0.50 | 0.27 | 3 | 0.1890 | |
| **A561E** | 0.14 | 0.07 | 3 | 0.39 | 0.30 | 3 | 0.2340 | |
| **D110E** | 0.93 | 0.58 | 3 | 109.63 | 8.65 | 3 | 0.0001 | |
| **D110H** | 0.42 | 0.34 | 3 | 93.95 | 6.55 | 3 | 0.0001 | |
| **D1152H** | 1.13 | 0.49 | 3 | 77.59 | 3.07 | 3 | 8.14E-06 | |
| **D1270N** | 1.55 | 0.32 | 3 | 185.46 | 19.12 | 3 | 0.0003 | |
| **D579G** | 0.09 | 0.09 | 3 | 43.30 | 14.86 | 3 | 0.0220 | |
| **E193K** | 0.62 | 0.26 | 3 | 42.22 | 9.76 | 3 | 0.0254* | |
| **E56K** | 0.25 | 0.25 | 3 | 18.26 | 16.01 | 3 | 0.1620 | |
| **E92K** | 0.14 | 0.14 | 3 | 0.32 | 0.24 | 3 | 0.2880 | |
| **F1052V** | 1.82 | 0.63 | 3 | 171.92 | 10.39 | 3 | 4.11E-05 | |
| **F1074L** | 0.98 | 0.44 | 3 | 69.19 | 6.21 | 3 | 0.0041* | |
| **F508del** | 0.13 | 0.13 | 3 | 0.80 | 0.46 | 3 | 0.1150 | |
| **G1244E** | 0.43 | 0.33 | 3 | 7.20 | 6.21 | 3 | 0.1690 | |
| **G1349D** | 0.09 | 0.09 | 3 | 6.49 | 0.70 | 3 | 0.0004 | |
| **G178R** | 0.81 | 0.41 | 3 | 10.73 | 6.06 | 3 | 0.0890 | |
| **G551D** | 0.44 | 0.26 | 3 | 2.18 | 1.12 | 3 | 0.1349* | |
| **G551S** | 0.42 | 0.21 | 3 | 14.73 | 3.82 | 3 | 0.0100 | |
| **G85E** | 0.63 | 0.63 | 3 | 0.32 | 0.16 | 3 | 0.3270 | |
| **G970R** | 0.44 | 0.24 | 3 | 17.45 | 7.30 | 3 | 0.0405 | |
| **H1054D** | 0.26 | 0.26 | 3 | 1.89 | 0.17 | 3 | 0.0030 | |
| **H1085R** | 0.26 | 0.24 | 3 | 1.08 | 0.37 | 3 | 0.0685 | |
| **I336K** | 0.95 | 0.22 | 3 | 43.76 | 12.94 | 3 | 0.0150 | |
| **I507del** | 0.20 | 0.20 | 3 | 0.85 | 0.29 | 3 | 0.0680 | |
| **K1060T** | 2.24 | 1.44 | 3 | 89.88 | 4.34 | 3 | 2.18E-05 | |
| **L1065P** | 0.15 | 0.15 | 3 | 0.37 | 0.19 | 3 | 0.2100 | |
| **L1077P** | 0.30 | 0.30 | 3 | 0.17 | 0.11 | 3 | 0.3475 | |

| | | | | | | | | |
|---|---|---|---|---|---|---|---|---|
| **L206W** | 0.48 | 0.22 | 4 | 12.35 | 9.55 | 3 | 0.0990 | |
| **L467P** | 0.04 | 0.04 | 3 | 0.58 | 0.39 | 3 | 0.1200 | |
| **L927P** | 0.04 | 0.04 | 3 | 1.26 | 0.56 | 3 | 0.0799* | |
| **M1101K** | 0.12 | 0.06 | 3 | 2.41 | 0.46 | 3 | 0.0040 | |
| **N1303K** | 0.16 | 0.12 | 3 | 0.45 | 0.13 | 3 | 0.0905 | |
| **P67L** | 0.13 | 0.12 | 3 | 2.23 | 1.37 | 3 | 0.1010 | |
| **R1066C** | 0.03 | 0.02 | 3 | 0.60 | 0.25 | 3 | 0.0435 | |
| **R1066H** | 0.85 | 0.37 | 3 | 7.92 | 5.38 | 4 | | 0.057 |
| **R1066M** | 0.05 | 0.05 | 3 | 0.67 | 0.32 | 3 | 0.0650 | |
| **R1070Q** | 3.39 | 0.76 | 4 | 197.94 | 9.85 | 4 | 0.0001* | |
| **R1070W** | 0.54 | 0.30 | 4 | 40.38 | 5.56 | 4 | | 0.029 |
| **R117C** | 0.27 | 0.13 | 3 | 40.97 | 11.27 | 3 | 0.0115 | |
| **R117H** | 0.75 | 0.36 | 3 | 46.27 | 12.73 | 3 | 0.0351* | |
| **R1283M** | 0.35 | 0.09 | 3 | 19.96 | 5.88 | 3 | 0.0145 | |
| **R334W** | 0.82 | 0.69 | 3 | 28.00 | 9.49 | 3 | 0.0230 | |
| **R347H** | 0.31 | 0.20 | 3 | 40.22 | 6.14 | 3 | 0.0015 | |
| **R347P** | 0.18 | 0.10 | 3 | 1.11 | 0.57 | 3 | 0.0915 | |
| **R352Q** | 0.30 | 0.12 | 3 | 25.83 | 5.88 | 3 | 0.0060 | |
| **R560S** | 0.04 | 0.04 | 3 | 0.31 | 0.21 | 3 | 0.1395 | |
| **R560T** | 0.24 | 0.21 | 3 | 0.70 | 0.43 | 3 | 0.1950 | |
| **R74W** | 1.13 | 0.39 | 3 | 178.21 | 16.56 | 3 | 0.0002 | |
| **S1251N** | 0.65 | 0.20 | 3 | 41.08 | 7.06 | 3 | 0.0025 | |
| **S1255P** | 0.21 | 0.19 | 3 | 22.07 | 6.06 | 3 | 0.0115 | |
| **S341P** | 0.28 | 0.28 | 3 | 8.80 | 4.85 | 3 | 0.0770 | |
| **S492F** | 0.30 | 0.23 | 3 | 0.69 | 0.48 | 3 | 0.2465 | |
| **S549N** | 0.25 | 0.12 | 3 | 11.00 | 8.37 | 3 | 0.1340 | |
| **S549R** | 0.25 | 0.25 | 3 | 4.35 | 4.09 | 3 | 0.1870 | |
| **S945L** | 0.00 | 0.00 | 3 | 17.70 | 9.55 | 3 | 0.1024* | |
| **S977F** | 0.45 | 0.10 | 3 | 121.72 | 15.37 | 3 | 0.0005 | |
| **T338I** | 0.25 | 0.15 | 4 | 27.53 | 7.35 | 4 | 0.0050 | |
| **V520F** | 0.06 | 0.06 | 3 | 1.19 | 0.86 | 3 | 0.1290 | |
| **Y569D** | 0.10 | 0.09 | 3 | 1.00 | 0.40 | 3 | 0.0460 | |
| **WT** | **2.53** | **0.54** | **7** | **180.31** | **15.97** | **6** | **0.0001*** | |

**Supporting Table S7**

Comparison of CFTR conductance ($G_{CFTR}$_norm, nS) after baseline activation with 10 μM forskolin (**fsk**) vs. potentiated activation with 10 μM forskolin + 10 μM VX-770 (a) (**fsk + VX-770**). Statistical significance of difference was assessed as described for table S6. In this dataset, normality test failed for G85E, R1066C, R1066H and S549R.

| Mutation | fsk | SEM | n | fsk+ VX-770 | SEM | n | P value (t test) | P value (Rank Sum test) |
|---|---|---|---|---|---|---|---|---|
| **A455E** | 0.88 | 0.33 | 3 | 5.46 | 0.59 | 3 | 0.0005 | |
| **A46D** | 2.14 | 1.16 | 3 | 19.02 | 1.34 | 3 | 0.0002 | |
| **A559T** | 0.50 | 0.27 | 3 | 0.74 | 0.15 | 3 | 0.1215 | |
| **A561E** | 0.39 | 0.30 | 3 | 0.31 | 0.11 | 4 | 0.2018 | |
| **D110E** | 109.63 | 8.65 | 3 | 214.38 | 3.61 | 3 | 0.0001 | |
| **D110H** | 93.95 | 6.55 | 3 | 205.40 | 6.63 | 3 | 0.0001 | |
| **D1152H** | 77.59 | 3.07 | 3 | 213.55 | 13.93 | 3 | 0.0002 | |
| **D1270N** | 185.46 | 19.12 | 3 | 207.96 | 13.18 | 3 | 0.0970 | |
| **D579G** | 43.30 | 14.86 | 3 | 140.17 | 25.68 | 3 | 0.0078 | |
| **E193K** | 42.22 | 9.76 | 3 | 243.15 | 15.27 | 3 | 0.0001* | |
| **E56K** | 18.26 | 16.01 | 3 | 38.96 | 8.62 | 3 | 0.0795 | |
| **E92K** | 0.32 | 0.24 | 3 | 0.79 | 0.28 | 3 | 0.0675 | |
| **F1052V** | 171.92 | 10.39 | 3 | 213.18 | 4.42 | 3 | 0.0055 | |
| **F1074L** | 69.19 | 6.21 | 3 | 152.94 | 5.98 | 3 | 0.0001 | |
| **F508del** | 0.80 | 0.46 | 3 | 4.29 | 0.80 | 3 | 0.0035 | |
| **G1244E** | 7.20 | 6.21 | 3 | 50.71 | 15.47 | 3 | 0.0148 | |
| **G1349D** | 6.49 | 0.70 | 3 | 202.73 | 5.63 | 3 | 1.03E-06 | |
| **G178R** | 10.73 | 6.06 | 3 | 232.07 | 25.17 | 3 | 0.0003 | |
| **G551D** | 2.18 | 1.12 | 3 | 35.13 | 7.56 | 3 | 0.0033 | |
| **G551S** | 14.73 | 3.82 | 3 | 207.71 | 35.87 | 3 | 0.0015 | |
| **G85E** | 0.32 | 0.16 | 3 | 0.31 | 0.16 | 3 | | 1 |
| **G970R** | 17.45 | 7.30 | 3 | 183.91 | 7.78 | 3 | 1.63E-05 | |
| **H1054D** | 1.89 | 0.17 | 3 | 40.92 | 6.29 | 3 | 0.0008 | |
| **H1085R** | 1.08 | 0.37 | 3 | 28.72 | 5.22 | 3 | 0.0015 | |
| **I336K** | 43.76 | 12.94 | 3 | 139.96 | 20.78 | 3 | 0.0043 | |
| **I507del** | 0.85 | 0.29 | 3 | 0.36 | 0.36 | 3 | 0.0858 | |
| **K1060T** | 89.88 | 4.34 | 3 | 197.15 | 7.63 | 3 | 3.55E-05 | |
| **L1065P** | 0.37 | 0.19 | 3 | 2.07 | 0.79 | 3 | 0.0260 | |
| **L1077P** | 0.17 | 0.11 | 3 | 0.27 | 0.21 | 3 | 0.1715 | |
| **L206W** | 12.35 | 9.55 | 3 | 42.40 | 5.37 | 4 | 0.0080 | |
| **L467P** | 0.58 | 0.39 | 3 | 0.23 | 0.05 | 3 | 0.1055 | |
| **L927P** | 1.26 | 0.56 | 3 | 101.26 | 7.07 | 3 | 3.66E-05 | |

| | | | | | | | | |
|---|---|---|---|---|---|---|---|---|
| **M1101K** | 2.41 | 0.46 | 3 | 8.26 | 3.20 | 3 | 0.0539* | |
| **N1303K** | 0.45 | 0.13 | 3 | 14.93 | 2.59 | 3 | 0.0076* | |
| **P67L** | 2.23 | 1.37 | 3 | 20.65 | 5.93 | 3 | 0.0098 | |
| **R1066C** | 0.60 | 0.25 | 3 | 1.21 | 0.30 | 3 | | 0.4 |
| **R1066H** | 7.92 | 5.38 | 4 | 33.71 | 5.59 | 3 | | 0.057 |
| **R1066M** | 0.67 | 0.32 | 3 | 1.31 | 0.21 | 3 | 0.0415 | |
| **R1070Q** | 197.94 | 9.85 | 4 | 214.84 | 2.44 | 3 | 0.0533 | |
| **R1070W** | 40.38 | 5.56 | 4 | 160.94 | 14.03 | 3 | 0.0001 | |
| **R117C** | 40.97 | 11.27 | 3 | 202.45 | 7.36 | 3 | 0.0001 | |
| **R117H** | 46.27 | 12.73 | 3 | 203.39 | 4.16 | 3 | 0.0001 | |
| **R1283M** | 19.96 | 5.88 | 3 | 86.07 | 10.10 | 3 | 0.0013 | |
| **R334W** | 28.00 | 9.49 | 3 | 157.23 | 13.48 | 3 | 0.0003 | |
| **R347H** | 40.22 | 6.14 | 3 | 219.57 | 7.75 | 3 | 1.08E-05 | |
| **R347P** | 1.11 | 0.57 | 3 | 23.03 | 8.52 | 3 | 0.0155 | |
| **R352Q** | 25.83 | 5.88 | 3 | 222.40 | 5.71 | 3 | 3.28E-06 | |
| **R560S** | 0.31 | 0.21 | 3 | 0.45 | 0.05 | 3 | 0.1400 | |
| **R560T** | 0.70 | 0.43 | 3 | 0.01 | 0.01 | 3 | 0.0627* | |
| **R74W** | 178.21 | 16.56 | 3 | 193.08 | 1.97 | 3 | 0.1058 | |
| **S1251N** | 41.08 | 7.06 | 3 | 164.18 | 38.96 | 3 | 0.0090 | |
| **S1255P** | 22.07 | 6.06 | 3 | 221.11 | 9.61 | 3 | 1.37E-05 | |
| **S341P** | 8.80 | 4.85 | 3 | 42.08 | 13.12 | 3 | 0.0190 | |
| **S492F** | 0.69 | 0.48 | 3 | 9.38 | 7.39 | 3 | 0.0765 | |
| **S549N** | 11.00 | 8.37 | 3 | 193.79 | 10.57 | 3 | 3.51E-05 | |
| **S549R** | 4.35 | 4.09 | 3 | 51.69 | 4.75 | 3 | | 0.1 |
| **S945L** | 17.70 | 9.55 | 3 | 125.69 | 8.66 | 3 | 0.0003 | |
| **S977F** | 121.72 | 15.37 | 3 | 204.79 | 10.83 | 3 | 0.0030 | |
| **T338I** | 27.53 | 7.35 | 4 | 84.76 | 6.72 | 3 | 0.0008 | |
| **V520F** | 1.19 | 0.86 | 3 | 0.48 | 0.29 | 3 | 0.1190 | |
| **Y569D** | 1.00 | 0.40 | 3 | 0.20 | 0.11 | 3 | 0.0310 | |
| **WT** | **180.31** | **15.97** | **6** | **218.00** | **33.27** | **4** | **0.0715** | |

**Supporting Figure S8**

Rare mutation panel characterization – comparison between results obtained in this study and published datasets. Comparison between ion channel function vs. short-circuit Ussing Chamber measurements (**A, C, E, G**) and membrane density vs. immunoblot studies (**B, D, F, H**). (**A-D**) Measurements obtained in this study are compared to those reported in Yu et al. J Cyst Fibros. 11: 237-45 (2012) [42] and Van Goor et al. J Cyst Fibros. 13, 29-36 (2014) [40]. (**E-H)**. Comparisons between this study vs. Sosnay et al. Nat Genet. 45, 1160-1167 (2013) [41]. Note that different sets of genotypes were included in the different studies, and individual genotypes are not shown in the same colour in Figures 4, S8 (A-D) and S8 (E-H). For display purposes, the two conductance and the $\rho$ axes are shown with logarithmic scaling. With these transformations, squared correlation coefficients obtained by simple linear regression ($r^2$) are: 0.61 (A), 0.53 (B), 0.58 (E) and 0.74 (F). Using untransformed measurements, the corresponding $r^2$values are: 0.61 (A), 0.49 (B), 0.68 (E) and 0.66 (F).

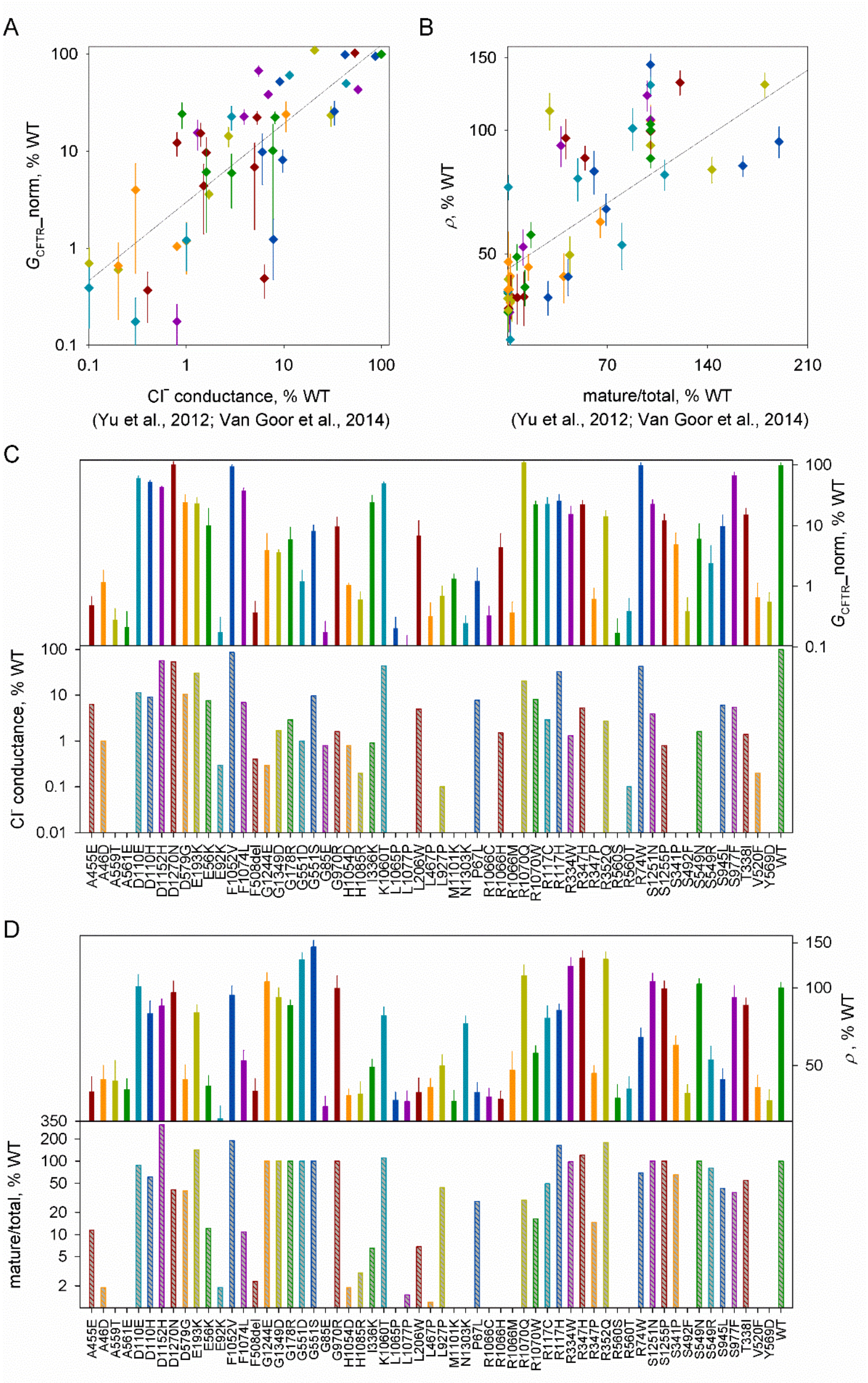
A
B
C
D
$G_{CFTR}$_norm, % WT
Cl⁻ conductance, % WT
(Yu et al., 2012; Van Goor et al., 2014)
ρ, % WT
mature/total, % WT
(Yu et al., 2012; Van Goor et al., 2014)
A455E A46D A559T A561E D110E D110H D1152H D1270N D579G E193K E56K E92K F1052V F1074L F508del G1244E G1349D G178R G551D G551S G85E G970R H1054D H1085R I336K K1060T L1065P L1077P L206W L467P L927P M1101K N1303K P67L R1066C R1066H R1066M R1070Q R1070W R117C R117H R334W R347H R347P R352Q R560S R560T R74W S1251N S1255P S341P S492F S549N S549R S945L S977F T338I V520F Y569D WT

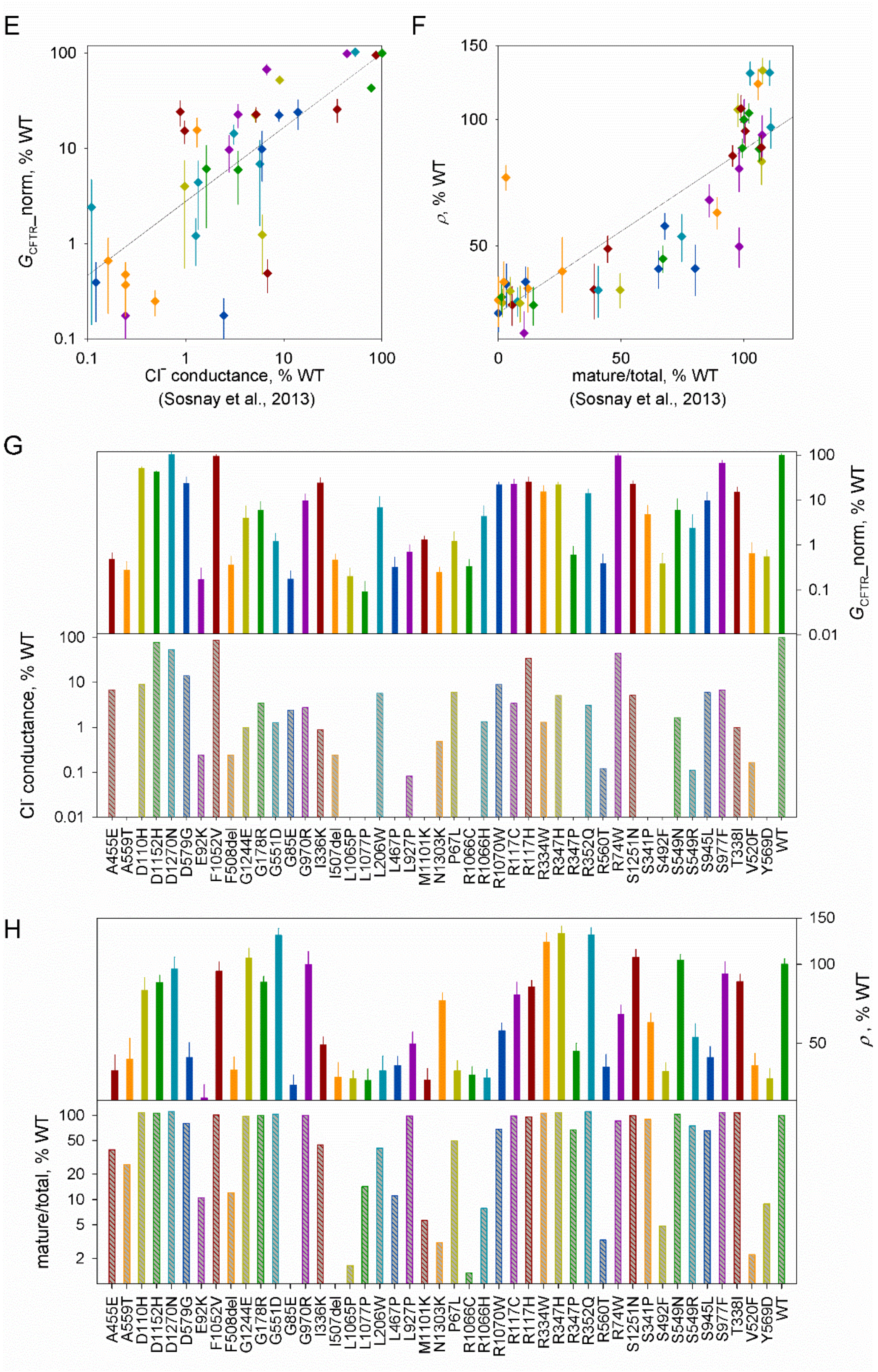
E
F
G
H
G_CFTR_norm, % WT
Cl⁻ conductance, % WT
(Sosnay et al., 2013)
ρ, % WT
mature/total, % WT
(Sosnay et al., 2013)
Cl⁻ conductance, % WT
G_CFTR_norm, % WT
mature/total, % WT
ρ, % WT
A455E A559T D110H D1152H D1270N D579G E92K F1052V F508del G1244E G178R G551D G85E G970R I336K I507del L1065P L1077P L206W L467P L927P M1101K N1303K P67L R1066C R1066H R1070W R117C R117H R334W R347H R347P R352Q R560T R74W S1251N S341P S492F S549N S549R S945L S977F T338I V520F Y569D WT

**Supporting Table S9**

Estimated values for fit parameters in traces for Figure 4B.

| Mutation | condition | $G_{CFTR}$ (nS) | $V_M$ (mV) | $G_{trans}$ (nS) | $\tau_{trans}$ (s) |
|---|---|---|---|---|---|
| **WT** | fsk | 152.88 | -59.42 | 14.8 | 5 |
| **S492F** | DMSO | 0.73 | -79.89 | 14.8 | 5 |
| | fsk | 1.603 | -69.19 | 14.8 | 5 |
| | fsk+VX-770 | 24.07 | -85.18 | 14.8 | 5 |
| **H1085R** | DMSO | 0.35 | -42.37 | 14.8 | 5 |
| | fsk | 1.28 | -51.01 | 19.1 | 3.94 |
| | fsk+VX-770 | 35.02 | -76.3 | 14.8 | 5 |
| **H1054D** | DMSO | 0.000125 | -58.76 | 14.8 | 5 |
| | fsk | 2.29 | -50.85 | 14.8 | 5 |
| | fsk+VX-770 | 30.17 | -70.69 | 14.8 | 5 |
| **L927P** | DMSO | 0.0000259 | -85.1 | 14.8 | 5 |
| | fsk | 1.47 | -41.2 | 14.8 | 5 |
| | fsk+VX-770 | 103.2 | -60.49 | 14.8 | 5 |
| **R1283M** | DMSO | 0.57 | -75.0 | 17.4 | 4 |
| | fsk | 25.86 | -75.3 | 14.8 | 5 |
| | fsk+VX-770 | 70.65 | -50.4 | 14.8 | 5 |

**Supporting Table S10**

Distances measured between α-carbons of highly VX-770 sensitive mutation sites and charged residues in the vicinity.

| Mutation | location | interacting charge | location | distance(Å) |
|---|---|---|---|---|
| **G178R** | TM3, ICL1 | K254<br>E257<br>R258 | TM4, ICL2 | 9.876<br>6.657<br>7.913 |
| **H1045D** | ICL4 | E543 | NBD1, X-loop | 8.875 |
| **H1085R** | TM11 | R1048 | TM10 | 6.277 |
| **N1303K** | NBD2 | R1358 | NBD2 | 7.843 |
| **G1349D** | NBD2 | γ-phosphate | ATP, site 1 | 6.173 |
